# Stall force measurement of the kinesin-3 motor KIF1A using a programmable DNA origami nanospring

**DOI:** 10.1101/2025.07.02.662715

**Authors:** Nobumichi Takamatsu, Hiroko Furumoto, Takayuki Ariga, Mitsuhiro Iwaki, Kumiko Hayashi

**Author notes:** Correspondence to Kumiko Hayashi and Mitsuhiro Iwaki.

## Abstract

DNA origami technology is a method for designing and constructing nanoscale structures using DNA, and it is being applied across various fields. This technology was advanced by developing the nanospring (NS), a fluorescently visible molecular spring that quantifies forces through its extension and has been used to measure myosin-generated forces. This study aims to measure the force exerted by the kinesin-3 motor protein KIF1A, mutations of which cause KIF1A-associated neurological disorder (KAND) and are associated with reduced force and motility. Unlike kinesin-1, KIF1A detaches easily under perpendicular loads, which can occur in optical tweezers experiments. By applying force parallel to the microtubule using the NS, we were able to precisely measure the stall force even for KAND mutants, for which such measurements are typically challenging. This result highlights the potential of the NS as a new tool for force spectroscopy in biophysics.

## Introduction

The kinesin superfamily of microtubule-based motor proteins in humans consists of approximately 45 types and represents a large group of motor proteins with diverse functions^1, 2^. Among them, KIF1A, a member of the kinesin-3 subfamily, plays a critical role in the long-distance transport of synaptic vesicle precursors and other cargos in neurons^3^. This transport is essential for maintaining synaptic activity, supporting neuronal development, and ensuring proper neuronal function. Structurally, KIF1A contains a motor domain that binds to microtubules and hydrolyzes ATP to generate mechanical force, driving movement toward the plus end of microtubules. In recent years, KIF1A has gained significant attention due to the identification of over 100 mutations linked to KIF1A-associated neurological disorder (KAND), a spectrum of neurodevelopmental conditions characterized by spastic paraplegia, intellectual disability, and progressive neurodegeneration^4, 5^.

Single-molecule measurements of KIF1A mutants associated with KAND have developed^6, 7, 8, 9^. Precise measurements of the velocity of KIF1A single molecules have been achieved using TIRF microscopy, while optical tweezers have enabled force measurements. These techniques have been used to compare the biophysical properties of KIF1A single molecules with those of its mutants that cause KAND. The motility of the single molecules has been reported to correlate with disease severity, highlighting the importance of single-molecule biophysical measurements into precision medicine^10^. The velocity of KIF1A has been extensively studied; however, there is still room for improvement in force measurements using optical tweezers. To date, optical tweezers have been used to measure the stall force, defined as the maximum force that a motor can generate when its movement is halted, of various microtubule-based motor proteins^11, 12, 13, 14, 15, 16, 17^. Since a submicron-sized bead is attached to the kinesin (∼10nm) and force is applied via the bead to the kinesin in this method, the applied force tends to pull it upward, which can sometimes lead to its detachment from the microtubule^18, 19^. Since KIF1A is easier to detach from microtubules compared to conventional kinesin-1, it is more challenging to successfully stall KIF1A in optical tweezer experiments without detachment. Indeed, most of the stalls lasted less than one second, and the force trajectories exhibited sawtooth-like patterns in previous studies^6, 7^.

To overcome the challenges associated with force measurements in KAND mutants, we employed an alternative approach based on DNA nanotechnology. In a previous study, Iwaki et al. developed a distinct method for force measurement by using programmable DNA origami to construct nanoscale, coil-like springs known as nanosprings (NSs)^20, 21^. These NSs are fluorescently labeled, allowing their extension to be visualized with conventional fluorescence microscopy. Since the force–extension relationship of the NS is pre-calibrated using acoustic force spectroscopy^21^, the force exerted on the spring can be quantitatively estimated from its measured extension. In this study, we applied the NS method to measure the force generated by single KIF1A molecules, including both wild-type and KAND mutants. Using the NS system, we were able to apply a horizontal load parallel to the microtubule axis to single KIF1A molecules, and clearly observed sustained stalling for several tens of seconds with reduced detachment. Notably, stalling events were also stably observed in KAND mutants such as P305L and V8M, which disrupt critical force-generating regions of the protein (e.g., the K-loop and neck linker) ^6, 7^. We also included the A255V mutant, previously unmeasured in force assays, which is known to affect ATP hydrolysis^6^. Furthermore, the NS allowed for precise stall force measurement, revealing force differences between KAND mutant homodimers and heterodimers composed of a wild-type and the mutant subunit. These results highlight the advantage of our approach in minimizing the force component perpendicular to the microtubule’s long axis.

In previous single-molecule experiments in kinesin, DNA origami was used to control the number of cooperatively moving kinesin motors, and their velocities have been analyzed^22, 23^. As a distinct application of DNA-origami-based motor protein systems, in this study, we successfully measured force using only fluorescence imaging. This study is significant in that it extends fluorescence imaging from merely capturing velocity to also allowing the quantification of force. In the field of motor proteins research, measurements of physical quantities related to distance, such as velocity and stepping motion, have rapidly advanced with the development of cutting-edge microscopy techniques like a super-resolution microscopy technology MINFLUX^24^, enabling their investigation not only in glass chambers but also inside living cells^25^. However, since both force and velocity are crucial physical parameters to understand the functions of motor proteins, and force-velocity relationships are indeed related to their ATP hydrolysis mechanisms^11, 26^, we believe that the development of a new type of force measurement is of great significance in this field. The development of NS-based force measurement, which enables force quantification at the nanoscale, is expected to have significant implications not only in the field of motor protein research but also in the broader study of protein biophysics.

## Results

### The NS-KIF1A-inert KIF5B complex as a force measurement system

To measure the force generated by single KIF1A molecule, we constructed a complex comprising a KIF1A molecule, an inert KIF5B mutant, and a NS. For KIF1A, the motor domain (amino acid residues 1–393) was used, excluding the C-terminal and cargo-binding domains (Fig. 1a). Since KIF1A does not form stable dimers in the absence of cargo binding domains, a leucine zipper domain was incorporated to stabilize KIF1A dimers, mimicked the activated state of full-length KIF1A, as in the previous study ^6^. As for the inert KIF5B attached to the one end of the NS, it was engineered to remain immobilized on microtubules by introducing the G234A mutation (Methods). This mutant lacks ATP hydrolysis activity and, as a result, does not exhibit motility, allowing it to remain stably attached to microtubules ^27^. Both kinesins were equipped with SNAP-tags at their N-termini, which enabled chemical coupling to the NS (Fig. 1b; Methods). The inert KIF5B mutant served to anchor the KIF1A–NS complex to a microtubule. This anchoring of the KIF5B to the microtubule allowed a KIF1A to stay near the microtubule, increasing the probability of its binding to the microtubule.

**Fig. 1.**
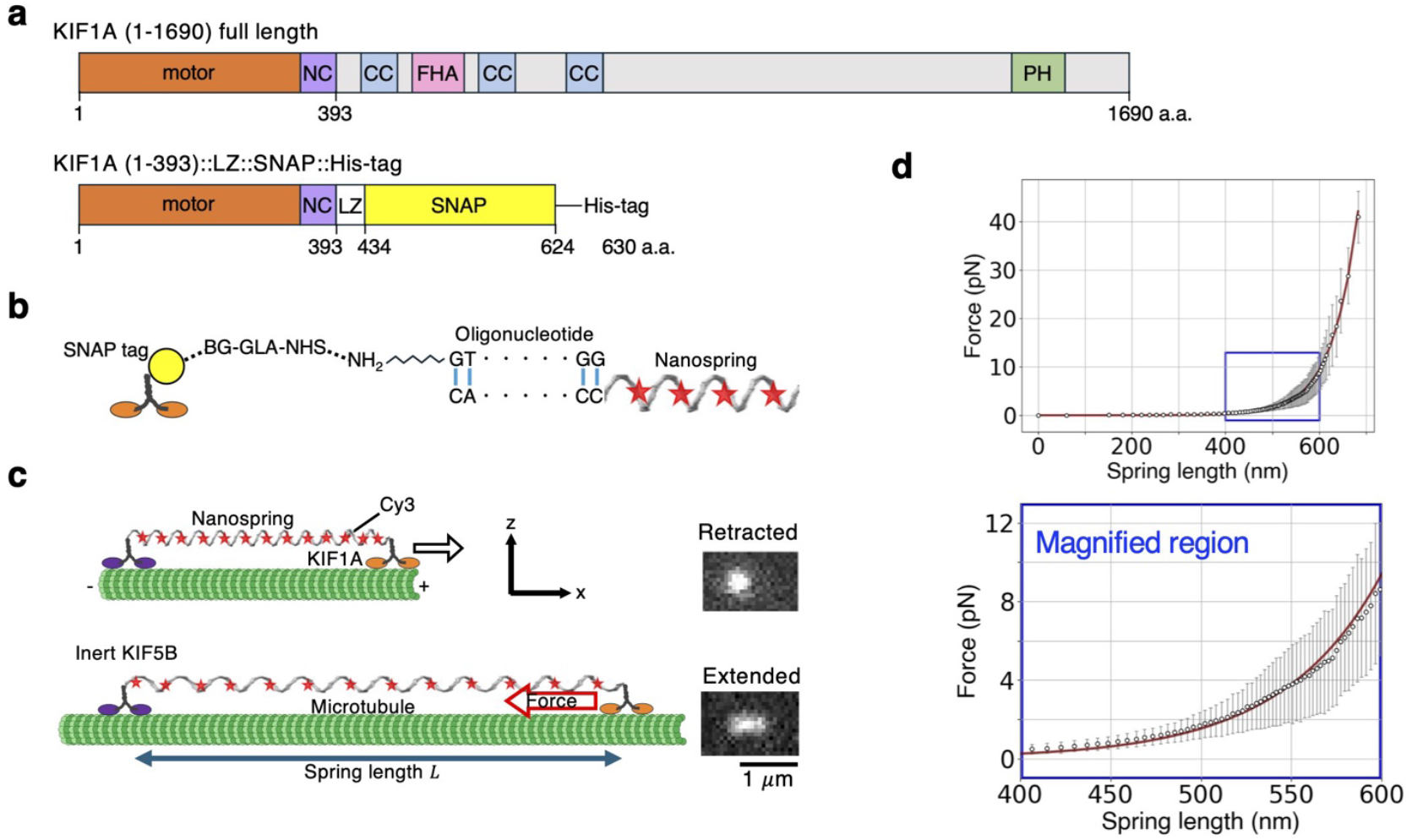
Experimental design for the stall force measurement of KIF1A using a NS. **a** Schematic of the domain structure of full-length KIF1A and the recombinant construct used in the experiments. To stabilize KIF1A dimers, which do not form stably without cargo-binding domains, a leucine zipper was incorporated, and SNAP-tags were added at the N-termini to enable chemical coupling to the NS. **b** Schematic of the chemical modifications required for coupling the NS to kinesin (Methods). **c** An inert KIF5B is anchored to the microtubule, and the NS extends as a KIF1A moves toward the plus end. The NS is uniformly labeled with Cy3 fluorophores, allowing force to be calculated from its extension. The micrographs depict the NS in the retracted and extended states. Here, the microtubule axis is defined as the *x*-direction, and the direction perpendicular to the microtubule is defined as the *z*-direction. **d** Force–extension relationship of the NS, showing nonlinear elastic behavior, calibrated by acoustic force spectroscopy (AFS)^21^ and fitted with an exponential function (Methods).

At equilibrium, in the absence of tension applied by KIF1A, the NS, uniformly labeled with 124 Cy3 fluorophores, exhibited a resting length of approximately 300 nm and appeared as a diffraction-limited circular spot in a fluorescence image (Fig. 1c). In the presence of ATP, the NS extended as the KIF1A moved against the load, and its fluorescence image became an elongated shape (Fig. 1c). To calibrate the pulling force acting on the NS from its extension, the acoustic force spectroscopy (AFS) was used as in the previous study^21^. Figure 1d presents the relationship between NS extension and force measured by the AFS, revealing its nonlinear elastic behavior. Its non-linear force-extension relationship was fitted with an exponential function (Methods).

### Simulation-based evaluation of fluorescence image analysis methods

Before analyzing the fluorescence images of a NS, we generated artificial images that mimic experimental conditions in order to develop a method for estimating the length of a rod-shaped object from its fluorescence image. The artificial images were generated by placing 116 two-dimensional isotropic Gaussian functions aligned along a straight line (Fig. 2a), to mimic a DNA calibration rod labeled with 116 fluorescent molecules, as introduced in the following section. The value of the variance in the Gaussian function was determined based on the approximate shape of the point spread function (PSF) of the Cy3 fluorophore (Fig. 2b). While the PSF is, in principle, influenced by the optical system, here it was used as a reference value for generating the artificial images. White noise was then added to the images. When the 116 Gaussian functions were superimposed, they appear as a single bright spot with an elliptical shape (Fig. 2c, top). For this artificial image, we present a comparison of the results obtained using two methods: the Gaussian fitting method and the chain fitting method.

**Fig. 2.**
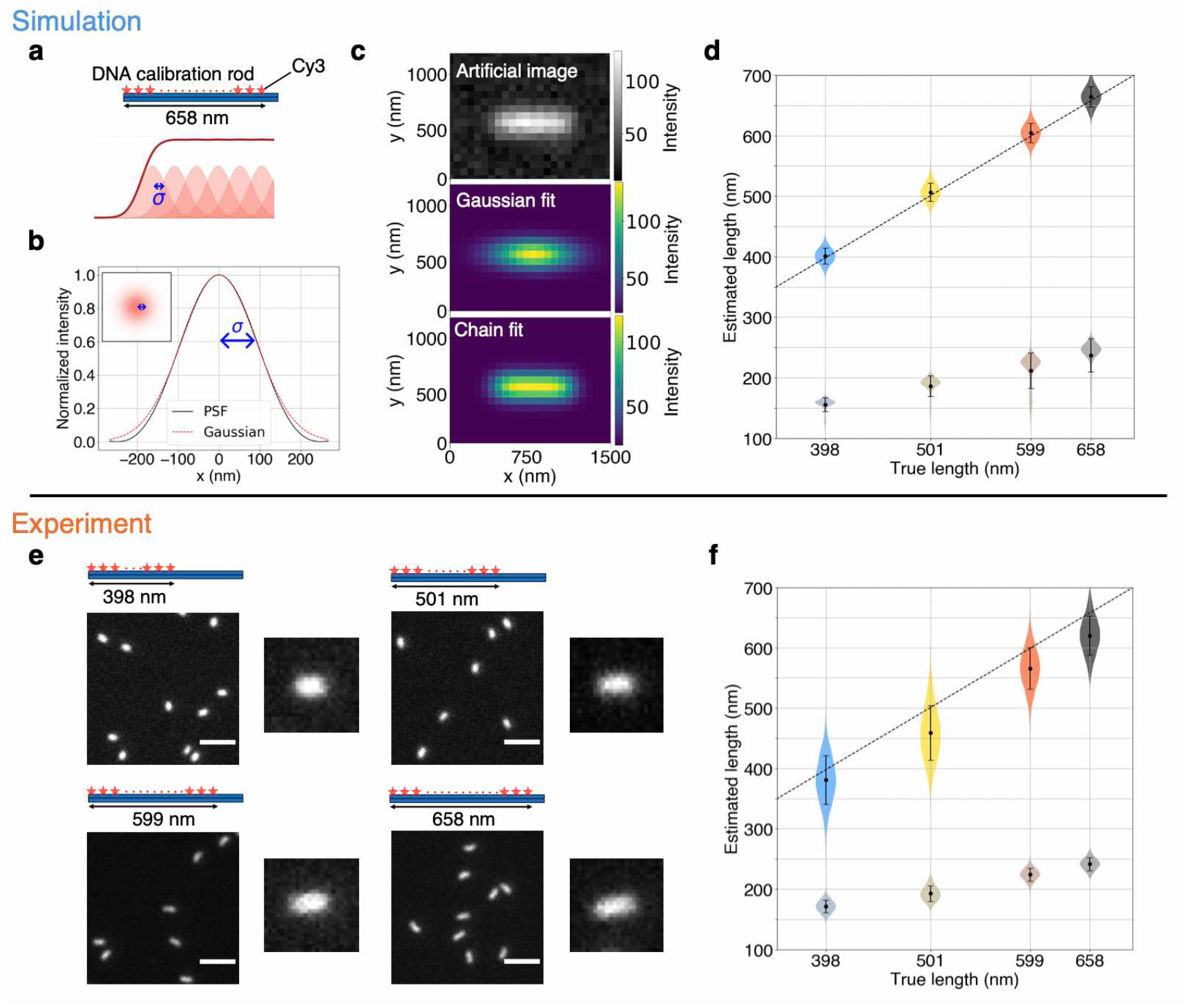
Length estimation using DNA calibration rods. **a** The fluorescence intensity of the DNA calibration rod was theoretically modeled as a superposition of Gaussian functions with variance *σ*^2^ aligned along a straight line, and an artificial image was generated accordingly. **b** The *σ* value of the Gaussian function used in the model was fitted to match the point spread function of Cy3 fluorescent dye (black line), resulting in a value of 90.65 nm. **c** Artificial image generated by placing 116 two-dimensional Gaussian functions (top). When 116 Gaussian functions were superimposed, they appeared as a single bright spot with an elliptical shape. The image (top) was fitted by the Gaussian fitting method (equation (1)) (middle) and the chain fitting method (equation (2)) (bottom). **d** Estimated length plotted against the true length of the model (*L*) using the Gaussian fitting method (equation (1)) (dark colors) and the chain fitting method (equation (2)) (bright colors), respectively. The dotted line represents the linear equation *y* = *x*. The chain fitting model provides estimates the true value of *L* in the simulation. **e** Fluorescence micrographs of the DNA calibration rods^21^ obtained in real experiments, with lengths of 398 nm, 501 nm, 599 nm, and 658 nm. The scale bars indicate 2 μm. **f** Estimated length plotted against the true length of the rods using the Gaussian fitting method (equation (1)) (dark colors) and the chain fitting method (equation (2)) (bright colors), respectively. The dotted line represents the linear equation *y* = *x*. The chain fitting method provides estimates closer to the true value of *L*.

In the Gaussian fitting method, the whole fluorescence spot is approximated using a single two-dimensional Gaussian function *I*_*g*_(*x, y*), and the estimated the standard deviation along the long axis (*σ*_long_) (Fig. 2c, middle).

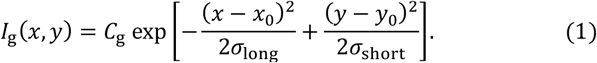

Here, *C*_*g*_ is a constant, and *x* and *y* represent the major and minor axis directions, respectively. Figure 2d (gray symbols) shows the relationship between the true length *L* of the rod shaped object and *σ*_long_. If the linear relationship between *L* and *σ*_long_ is determined in advance, the value of *L* can be estimated from the measured *σ*_long_.^21^

While the Gaussian fitting method provided reasonable results, we considered an alternative approach, called the chain fitting method, to directly estimate the length *L* of the model. In the Gaussian fitting method, proposed in the previous study^21^, the true length *L* of the rod is not included as a parameter in the fitting function (equation (1)). The chain fitting method is designed based on the structure in which fluorescent molecules are arranged in a line, and because the fitting parameters directly include the length *L*, it is expected to provide a more accurate estimation than the Gaussian fitting method. The fluorescence intensity of this model can be described by the following equation:

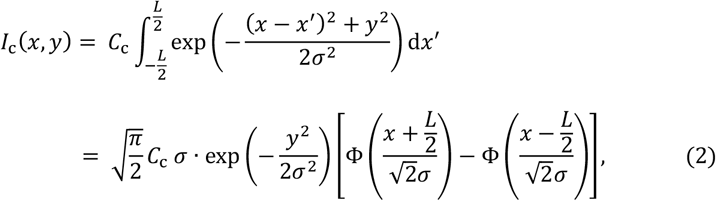

where *C*_c_ is a constant, and Φ is the error function. Figure 2c, bottom shows the resulting image obtained by the chain fitting method. The simulation results indeed demonstrated that the estimated length by the chain fitting method corresponded with the true length *L* (Fig. 2d, color symbols).

### Experimental validation of the chain fitting method using DNA calibration rods

We experimentally validated our approach of the chain fitting method using DNA calibration rods with known lengths.^21^ Figure 2e shows the fluorescence images of biotinylated DNA calibration rods immobilized on a glass surface via biotin-BSA and streptavidin (Methods), recorded under the same conditions used for observation of the NSs. As illustrated in the schematic, a total of 116 Cy3 molecules were conjugated to the DNA calibration rod at equal intervals ^21^. The length was varied by adjusting the interval between fluorescent dyes. The end-to-end distances between fluorescent molecules were designed to be 398 nm, 501 nm, 599 nm, and 658 nm, respectively. Since the NSs were labeled with 124 Cy3 fluorescent molecules, we considered the fluorescence images obtained from these rods to be similar to those of the NSs.

Figure 2f shows the *σ*_long_ and length *L* estimated from experimental fluorescence images of DNA calibration rods using the Gaussian fitting method (equation (1)) and chain fitting method (equation (2)), respectively. There is a proportional relationship between *σ*_long_ and the true length. On the other hand, *L* estimated by equation (2) provides a direct estimate close to the true length of the DNA calibration rod. However, the fact that *L* was approximately 10% shorter than the true length is considered to be likely due to slight bending of the DNA-origami structures caused by thermal fluctuations. Indeed, in the case of longer rods shown in Fig. 2e, a slight bending was observed. Accordingly, the length was corrected by a factor of 1.07 (Methods). In the following, the extension of the NS was primarily estimated using the chain fitting method. Note that results obtained using the conventional Gaussian fitting method are also presented in the Supplementary Information. The results from both methods were consistent.

### Observation of stall behavior in wild-type KIF1A

The supplementary movie shows a representative example of the movement of a NS with KIF1A and KIF5B. At first an inert KIF5B is firmly fixed to the microtubule. Then, the diffusive behavior of a KIF1A—attached to the NS, which is anchored to the microtubule via the inert KIF5B—as it searches for a binding site on the microtubule, is observed. Once the KIF1A binds to the microtubule, it starts moving along the microtubule and stretches the spring. When the tension in the NS balances with the maximum force generated by the KIF1A, the extension temporarily halts during the time interval (attachment duration). Eventually, the KIF1A detaches from the microtubule, the tension is released, and the NS returns to its retracted state. This cycle event was repeatedly observed.

The extension of the NS was estimated by the chain fitting method (equation (2)) (Fig. 3a, bottom). For the event during a time interval (2.8 s ≤ *t* ≤ 21.2 s), the histogram of the spring length was calculated (Fig. 3a, left). The histogram exhibits a bimodal Gaussian distribution, with each peak corresponding to the extended and retracted states of the NS shown in the micrographs (Fig. 3a left). *L*_stall_ was calculated by averaging the NS extension during the force-plateau periods within each attachment event. The attachment of KIF1A to the microtubule was identified by the marked suppression of angular fluctuations of the NS relative to the microtubule (Methods and Fig. 3a, top), which clearly distinguished the bound state from the diffusive search behavior. Within this attachment duration, the force plateau (stall duration Δ*t*) was detected by analyzing the rate of relative increase in NS’s length (Methods). The mean NS extension over these plateau segments was defined as *L*_stall_.

**Fig. 3.**
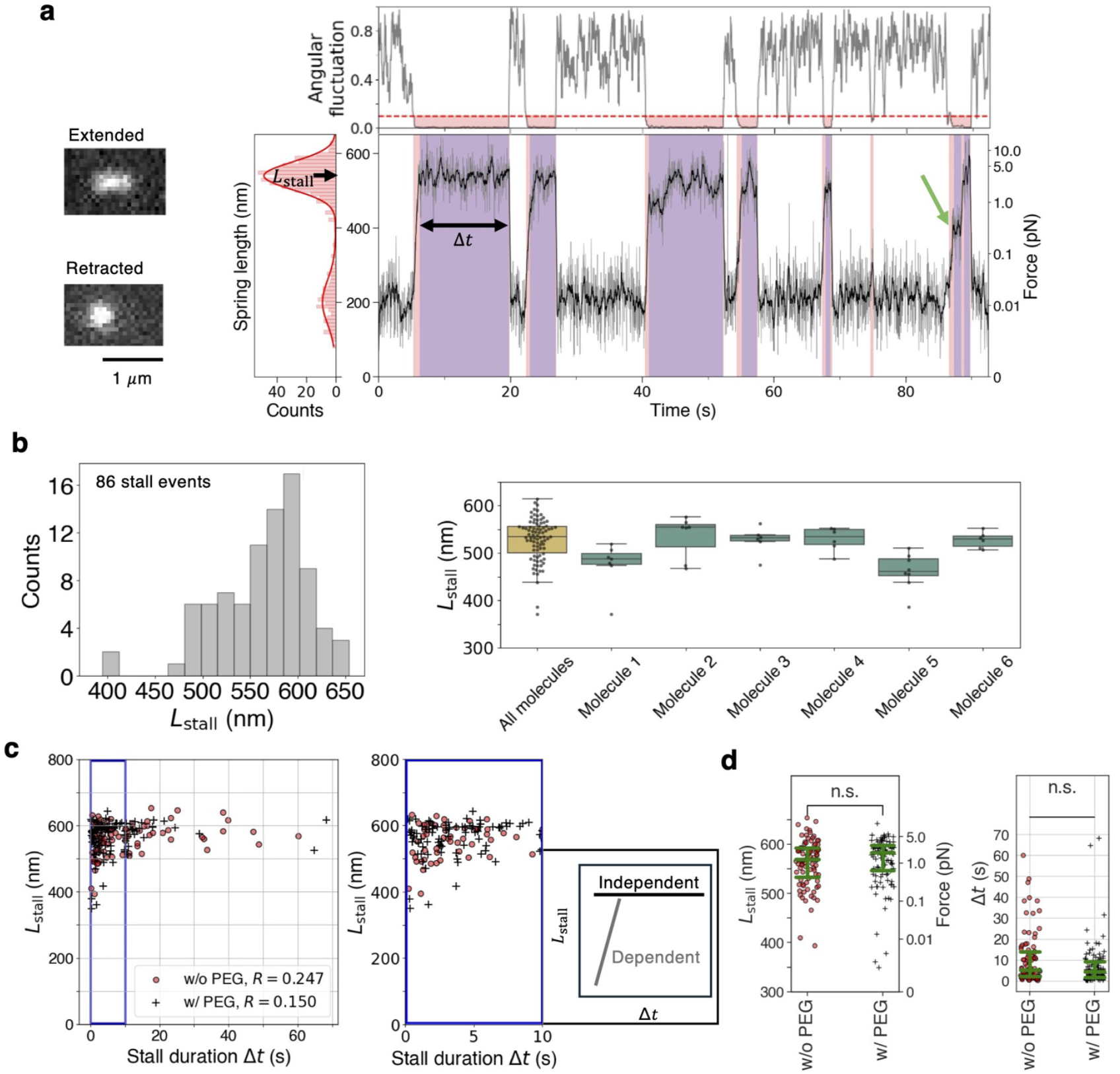
Stall force measurement of wild type KIF1A homodimers using NSs. **a** Time course of NS extension (*L*(*t*)) in the case of wild-type KIF1A. The black line (trace) represents the average over 10 frames. As illustrated in the schematic in Fig. 1c, the NS is stretched (micrograph, top) as a KIF1A moves toward the plus end of the microtubule. When the load reaches the maximum force that the KIF1A can generate, a stall is observed, followed by the detachment of the KIF1A from the microtubule. The NS then returns to its original retracted state (micrograph, bottom). The stall duration Δ*t* (violet region) was defined based on the angular fluctuation and the rate of relative increase in NS’s length (Methods), where the red regions represent the attachment durations decided based on the angular fluctuations. For each stall event, the histogram of NS extension exhibits a bimodal Gaussian distribution, with the higher peak corresponding to the stall length *L*_stall_ (left panel). **b** Histogram of *L*_stall_ values calculated from 86 stall events (left). The right panel shows the distribution of stall forces for each KIF1A molecule in which six or more stall events were observed. **c** *L*_stall_ as a function of stall duration Δ*t* with (n=105) and without (n=86) PEG. The correlation coefficient (*R*) between *L*_stall_ and Δ*t* is shown in the figure (left). The right panel presents the magnified view of the blue rectangle in the left panel, and clearly indicates that *R* is small. **d** Comparison of *L*_stall_ and Δ*t* with and without PEG (Mann–Whitney U test, *p* = 0.4718 for *L*_stall_, *p* = 0.0616 for Δ*t*). n.s., not significant (*p* ≥0.05). The green bars indicate the median values along with the first and third quartiles.

In our experiments, we analyzed a total of 86 events from 24 KIF1A molecules (Fig. 3b, left). From the mean value of *L*_stall_ (562 ± 48 nm), the stall force of the wild type KIF1A was estimated to be 4.7 pN (Table 1) by using the force-extension calibration of the NS (Fig. 1d). This value is similar to the stall force values of the wild type KIF1A(1-393) measured previously^19^, and larger than the detachment force values^6, 7^. The right panel of Fig. 3b shows the distribution of stall forces for each KIF1A molecule in which six or more stall events were observed. The broad distribution of stall forces arises not only from differences between molecules, but also from the variability of stall forces within a single molecule. The broad distribution is caused by the presence of step-back events (*e*.*g*., a green arrow in Fig. 3a); however, the chemical state underlying these step backs in KIF1A remains unknown.

**Table 1.** Stall force of KIF1A(1-393) estimated from the average value of *L*_stall_. The stall force values were calculated from the mean values of *L*_stall_ by using the force-extention relationship of the NS (Fig. 1d). The error of *L*_stall_ value represents the standard deviation (SD).

|  | Stall force (pN) | Detachment force (pN) | Termination force (pN) |
| --- | --- | --- | --- |
| WT/WT | 4.7 (n=86)<br>($L_{\text{stall}} = 562 \pm 48$ nm) | 2.7<sup>a</sup><br>2.2<sup>b</sup> | 4<sup>c</sup><br>6<sup>d</sup> |
| P305L/P305L | 0.2 (n=58)<br>($L_{\text{stall}} = 375 \pm 49$ nm) | 0.7<sup>b</sup> | - |
| P305L/WT | 0.3 (n=41)<br>($L_{\text{stall}} = 402 \pm 54$ nm) | - | - |
| V8M/V8M | 2.2 (n=95)<br>($L_{\text{stall}} = 516 \pm 57$ nm) | 1.9<sup>a</sup> | - |
| V8M/WT | 1.0 (n=61)<br>($L_{\text{stall}} = 469 \pm 27$ nm) | - | - |
| A255V/A255V | 3.0 (n=83)<br>($L_{\text{stall}} = 535 \pm 46$ nm) | - | - |
| A255V/WT | 2.5 (n=40)<br>($L_{\text{stall}} = 524 \pm 26$ nm) | - | - |

Detachment force and termination force measured by using optical tweezers.
a, reported in reference<sup>6</sup>; b, reported in reference<sup>7</sup>; c, reported in referece<sup>19</sup> for the single-bead assay; d, reported in referece<sup>19</sup> for the three-beads assay.

In the results presented here, the extension of the NS was estimated using the chain fitting method (equation (2)). For comparison, we also present the results obtained using the conventional Gaussian fitting method (equation (1)) in the Supplementary Information. The two methods provided consistent values, suggesting that future analyses can be performed without the need for prior calibration using DNA rods, which is required by the Gaussian fitting method. In addition, while direct velocity measurements were difficult in the NS-based experiments, we analyzed the behavior under low-load conditions by overlaying the time courses of 10 stall events (Supplementary Information).

### Stall duration Δ*t*

In previous studies^6, 7^, most of the stalls lasted less than one second, and the force trajectories exhibited sawtooth-like patterns, which was a characteristic feature of force generation by KIF1A unlike kinesin-1, which exhibits clear stalling behavior^28^. However, in our experiments, we observed clear stall states lasting for several tens of seconds, similar to the stalling behavior observed with kinesin-1 (Fig. 3a). To quantify this observation, we measured the stall durations (Δ*t*) and analyzed the relationship between *L*_stall_ and Δ*t* (Fig. 3c).

In a single bead optical tweezers experiment, it was reported that Δ*t* was correlated with the detachment force when the detachment of a KIF1A from the microtubule was likely accelerated by forces applied perpendicular to the microtubule’s long axis (*z*-axis depicted in the schematics of Fig. 1c), due to the contact between a single bead and a underlying microtubule^18^. By using the three-bead optical tweezers system^18, 19^, which was a modified version of the conventional optical trapping assay, to minimize the *z*-component of the force, it was found that Δ*t* did not depended on the detachment force (schematic in Fig. 3c)^18^. Our experimental results also indicated that Δ*t* was independent of *L*_stall_, as the correlation coefficient was 0.2 (Fig. 3c), suggesting that the *z*-component of the force was effectively minimized.

### Effect of electrostatic repulsion on force measurement

Because the DNA-based NS is negatively charged, we investigated the effect of its electrostatic repulsion with negatively charged microtubules. This repulsion could potentially promote the detachment of KIF1A from the microtubule, raising a concern for the measurement. To minimize repulsive interactions between the NS and the microtubule, we coated the NS with oligolysine conjugated to polyethylene glycol (PEG) to render the spring electrically neutral (Methods). Based on a statistical test, there was little difference in *L*_stall_ and stall duration (Δ*t*) between the conditions with and without PEG (Fig. 3c,d; Mann–Whitney U test, *p* = 0.4718 for *L*_stall_, *p* = 0.0616 for Δ*t*). These results suggest that the negative charge of the NS had little influence on the stall force measurements.

### Stall force measurement of the KIF1A disease-associated variant P305L

P305 of the motor domain of KIF1A is located adjacent to the positively charged K-loop insertion region of KIF1A (Fig. 4a). The positively charged K-loop interacts with the negatively charged C-terminal loop of tubulin in microtubules, promoting a high binding affinity to microtubules^15, 29^. Therefore, a mutation at P305 directly affects the interaction between KIF1A and microtubules. In other words, the P305L mutation could impair the function of the K-loop, leading to a reduced binding affinity to microtubules^7, 8^. It was reported that the velocity and run length were reduced by half, and the force decreased fourfold, indicating that the P305L mutation impaired the motility of the motor^7^. The decrease in force generation in the P305L mutant is a serious defect.

**Fig. 4.**
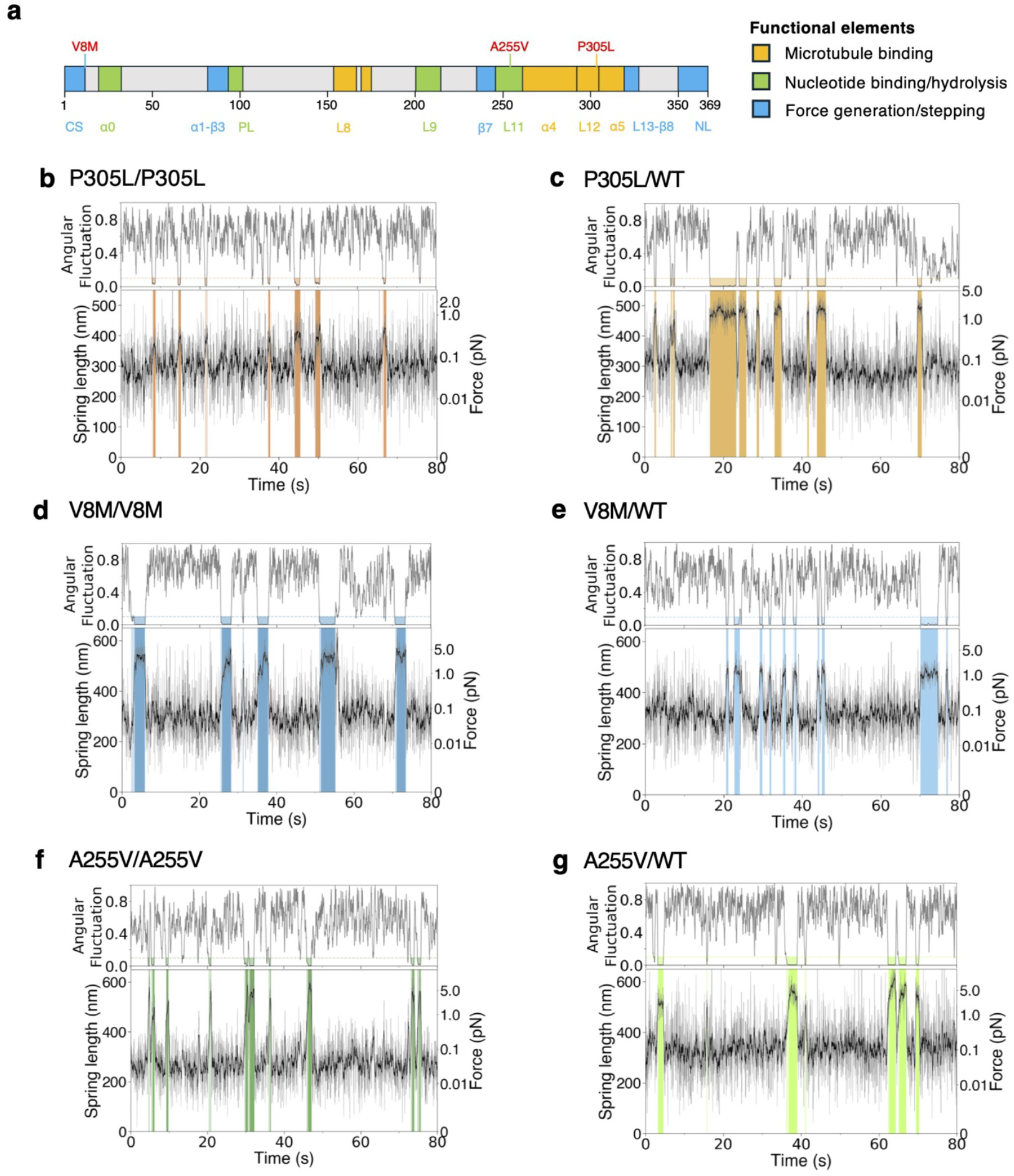
Stall force measurement of KAND mutants KIF1A homodimers and heterodimers using NSs. **a** Schematic of KIF1A domain structure showing the functions affected by the P305L, V8M, and A255V mutations^6^. Representative traces of NS extension are shown for homodimers and WT-mutant heterodimers of P305L, V8M, and A255V (**b**,**c**,**d**,**e**,**f**,**g**). The black lines (traces) represent the average over 10 frames. The lighter-colored regions in the graph represent the attachment durations, while the darker-colored regions indicate the stall durations. The identification of these durations is described in the Methods section.

The wild-type KIF1A depicted in Fig. 1b and 1c was replaced with the P305L mutant, and force measurements were performed using a NS for the mutant (Fig. 4b). Although it has been reported that the landing rate of P305L on microtubules was reduced by 97%^7, 8^, in this experiment, an inert KIF5B firmly anchored to the microtubule at the end of the NS, which kept the P305L mutant in close proximity to the microtubule, thereby effectively increasing its binding probability. In other words, the anchoring effect enabled clear observation of microtubule binding by the P305L mutant. This high binding rate is one of the advantages of the NS–KIF1A–inert KIF5B complex, when used as a force measurement system. Based on the value of *L*_stall_, the force generated by the mutant was reduced to be 0.2 pN (Table 1). Note that the histogram of *L*_stall_ is shown in Supplementary Information.

KIF1A is considered to primarily form dimers to anterogradely transport synaptic cargos in neurons. In recent years, the motility of heterodimers composed of wild-type and KAND-associated mutant subunits has begun to be investigated experimentally^6, 8, 30^. In heterozygous patients, WT/WT, P305L/WT, and P305L/P305L dimers coexist, highlighting the importance of studying the heterodimers. As in the previous study^8^, we constructed heterodimers composed of wild-type and P305L subunits (Methods) and performed stall force experiment using these heterodimers (Fig. 4c, Table 1).

### Stall force measurement of the KIF1A disease-associated variants V8M and A255V

Like kinesin-1, KIF1A also utilizes conformational changes in the neck linker to generate force^6^. The V8M mutation may hinder the neck linker from accessing its docking pocket, which in turn disrupts docking and impairs force generation (Fig. 4a). Using the NS-based stall force measurement, we also observed that the stall force of the V8M homodimer was reduced by approximately half (Fig. 4d, Table 1), consistent with the previous study^6^.

The A255 residue is located in loop L11 of the switch II cluster and may affect KIF1A function by altering the structure of L11 within switch II or the back door^31^ (Fig. 4a). In the single-molecule experiments, although its force has not yet been measured, a decrease in velocity has been reported^9, 32^. Loss of function in the switch II region or the back door may impair the conversion of ATP hydrolysis–driven conformational changes into mechanical force. Indeed, our NS-based stall force measurements revealed that the stall force was reduced by approximately half compared to the wild-type (Fig. 4f, Table 1). We also performed stall force measurements for heterodimers composed of the KAND mutants V8M (Fig. 4e, Table 1) or A255V (Fig. 4g, Table 1) and the wild-type subunit. Note that the histogram of *L*_stall_ for the mutants in this section is shown in Supplementary Information.

### *L*_stall_ values for different KIF1A variants

All *L*_stall_ values for the KIF1A mutants are summarized in Fig. 5a. The values increased in the order of P305L, V8M, and A255V. We then converted the *L*_stall_ values into forces and compared them with the clinical severity of KAND (Supplementary Information). Specifically, we compared them with REVEL, CADD v1.4, and the ESM score reported in the reference^5^. Among these metrics, the ensemble prediction score REVEL, which integrates multiple pathogenicity prediction algorithms, showed the correlation. It is important to expand the number of variants analyzed to further investigate the relationship between stall force and the severity of KAND.

**Fig. 5.**
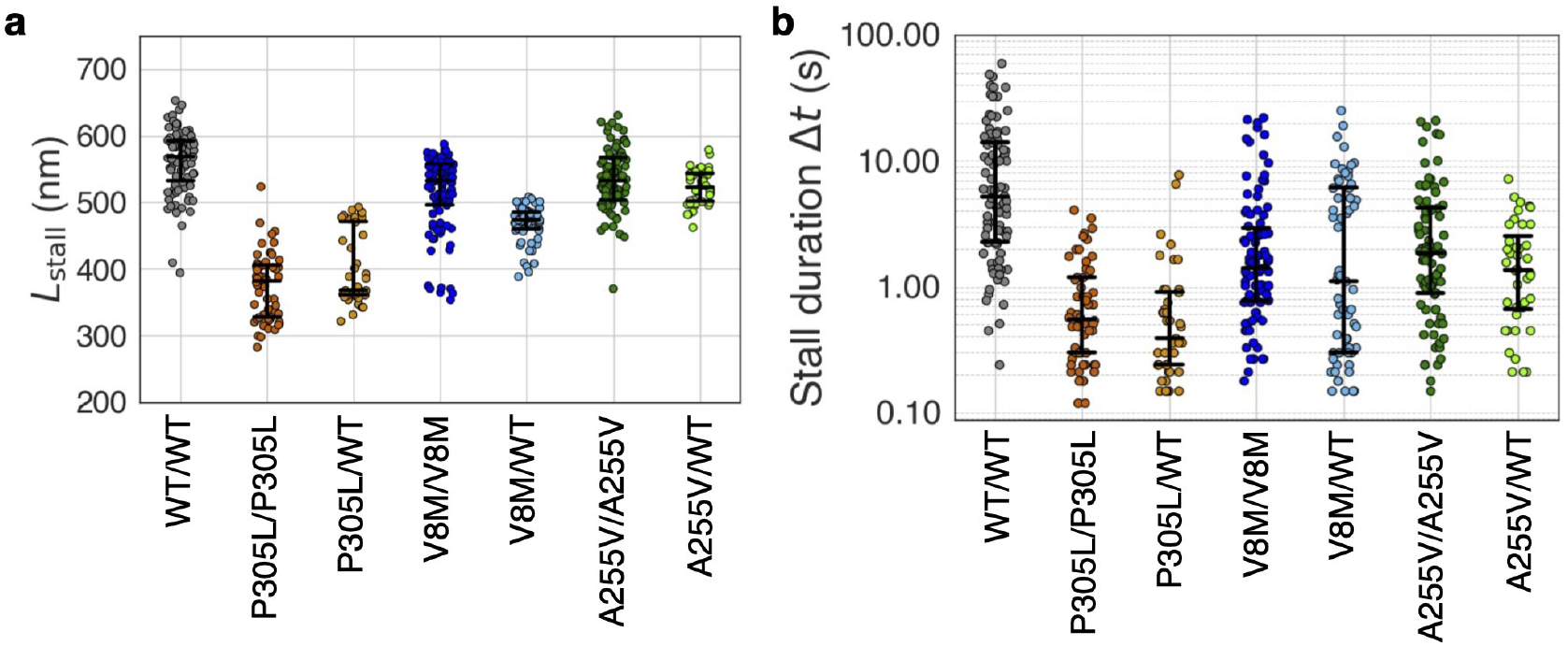
Comparison of *L*_stall_ and Δ*t* among KAND mutants. *L*_stall_ (**a**) and Δ*t* (**b**) for homodimers and heterodimers of P305L, V8M and A255V, compared with WT.

### Δ*t* values for different KIF1A variants

As shown in Fig. 5b, stall duration (Δ*t*) was compared among KAND mutants (P305L, V8M, and A255V). Because stall behavior has rarely been observed for KIF1A in single-bead optical tweezers experiments, comparison of Δ*t* has been limited in previous studies^6, 7^. Δ*t* did not show as clear differences among the mutants as *L*_stall_ did. For WT, which provided a sufficient number of stall events, we investigated the cumulative distribution of Δ***t*** (Supplementary Information). Understanding KIF1A’s behavior associated with these time constants will be important in future studies.

## Discussion

We measured the force generated by KIF1A using the NSs developed by Iwaki *et al*.^20, 21, 33^ (Fig. 1). Because KIF1A more readily detaches from microtubules compared to kinesin-1, and the *z*-component (refer to Fig. 1c for the z direction) of the applied forces often causes detachment in optical tweezers experiments, making it difficult to observe its stall behavior. In contrast, by using the NS, we were able to apply load to a single KIF1A molecule solely in the horizontal direction (*x*-axis) along microtubules, enabling clear observation of the stall events (Fig. 3). Note that, while performing these measurements, we also improved the image analysis method (Fig. 2). In a previous study^21^, the length of the NS was estimated using the Gaussian fitting method (equation (1)). In contrast, in the present study, we adopted the chain fitting method (equation (2)), which allowed for a more direct estimation of the length *L* from the images. Below, we discuss the advantages and future prospects of NS-based force measurements.

In KAND mutants, evolutionary scale model (ESM) scores have been compared with clinical phenotype scores such as the Vineland Adaptive Behavior Composite (VABS-ABC) scores ^34^. ESM refers to large-scale protein language models based on Transformer architectures, trained on millions of sequences to predict structure, function, and mutation effects from primary amino acid sequences^35^. However, due to the currently low correlation between ESM scores and clinical severity, incorporating biophysical parameters such as velocity and force may provide additional predictive insight, particularly for de novo mutations—a direction already explored in part by a previous study^10^. In the present study, we propose a high-throughput method for accurately measuring the force generated by KIF1A mutants using the NSs. Anchoring a KIF1A–NS complex to the microtubule via an inert KIF5B facilitated the observation of stall events even for KAND-related KIF1A mutants (Fig. 4), which were otherwise prone to detachment due to the dysfunctions caused by the mutations. Indeed, we successfully measured the stall force of several KAND-associated mutants, including P305L, V8M, and A255V. P305L, V8M, and A255V are representative mutants that primarily impair microtubule binding, force generation near the neck linker, and ATP hydrolysis near the Switch II region, respectively (Fig. 4a). Notably, stall force measurements for the homodimeric A255V and heterodimeric combinations P305L/WT, V8M/WT, and A255V/WT had not previously been reported using optical tweezers. Building a database of such biophysical measurements will be essential for linking molecular dysfunction to clinical outcomes. The NS-based force assays offer a powerful platform for this purpose. In the Supplementary Information, we additionally compared these three mutants with the REVEL, CADD v1.4, and ESM scores^5^. It is important to further investigate the relationship between stall force and the severity of KAND.

As noted in the introduction, optical tweezers have been the primary method for measuring forces generated by motor proteins^11, 12, 13, 14, 15, 36, 37, 38, 39^. An advantage of the NS is that it enables precise investigation of the force exerted by a single motor protein, whereas multiple kinesin molecules may attach to a bead in optical tweezers experiments. Moreover, unlike optical tweezers, which require individual manipulation of each bead and can measure the force of only one motor protein at a time, the NS-based assay allows simultaneous recording and force measurements of multiple NS–KIF1A complexes within the field of view. This results in significantly higher throughput and represents a key advantage over optical tweezers. From the perspective of compatibility with other measurement techniques, the near-infrared laser used in optical tweezers readily causes photobleaching of organic dyes, making single-molecule imaging of dye-labeled proteins challenging. For example, this means that optical tweezers cannot be used simultaneously with MINFLUX^25^. Moreover, the NS can also be combined with high-speed AFM or electron microscopy, enabling structural analysis of proteins under quantitatively controlled mechanical load. In fact, although not NSs, DNA origami-based thick filaments with attached myosin have been investigated using AFM^33^. Therefore, the DNA origami-based NSs are also expected to be used with AFM.

Although NSs offer several advantages, they also have disadvantages. While the optical trap can be approximated as a linear spring, the NS behaves as a nonlinear spring (Fig. 1d). Consequently, fluctuations in its length can sometimes lead to large errors in force measurements. Therefore, it is crucial to adjust the spring stiffness to match the force range generated by the motor protein. In this study, we used NSs with different physical properties from those used in the previous force measurements on myosin VI^11^. Wild-type KIF1A generates a larger stall force than myosin, requiring a different dynamic range for the nonlinear behavior of the spring. Owing to DNA origami technology, the mechanical properties of the NS can be tuned to match the appropriate force range for measuring molecular stiffness.

The reason why the stall force exhibits a wide range of values may be not only the non-linearity of the NS, but also the complexity of the underlying kinetics of a single KIF1A, such as step-back events (e.g., the green arrow in Fig. 3a). Indeed, analysis of the Δ*t* distribution (Fig. 6) suggests that the motion during stall events can be characterized by two (or more) reaction rate constants (Supplementary Information). Determining the detailed chemical reaction model underlying these stall events will be an important subject for future studies.

With the recent development of the super-resolution microscope MINFLUX, live-cell tracking of single fluorescent molecules has become possible, enabling the precise observation of kinesin-1 stepping motion in living cells with nanometer spatial and millisecond temporal resolution^25^. To date, models of chemo-mechanical energy conversion in motor proteins have been proposed based on the load dependence of their stepping motion ^26^. The intracellular environment is highly crowded with biomolecules and also contains active fluctuations^40^, resulting in complex viscoelastic properties and an inhomogeneous refractive index. This leads to challenges in using optical tweezers, such as accurately estimating the trap stiffness of laser-trapped vesicles in cells. On the other hand, the NSs offer a method of measuring force solely through fluorescence imaging, providing a potential solution to this problem. In relation to cellular applications, the NSs have already been used as force sensors to visualize the dynamics of integrins^21^. Expanding such applications, the introduction of the NS–kinesin complex into living cells offers a promising approach to uncover the load-dependent dynamics of kinesin in its native environment.

## Materials and Methods

### Plasmid Constructs

The plasmids used in this study were listed in Table S1 (Supplementary Information). A recombinant human KIF1A(1–393)-LZ-SNAP-6His construct containing the motor domain (amino acids 1–361), neck linker, neck coil domain, and a GCN4 leucine zipper (LZ) for stabilized dimerization was used for NS force generated study (Fig. 1a). To generate this construct, a human KIF1A(1–393)-LZ-6His construct named pSN672 (addgene #177362)^8^ was modified as follows: SNAP-tag cDNA was amplified by polymerase chain reaction (PCR) with high fidelity PCR enzyme (Prime STAR MAX, Takara Bio Inc., Kusatsu, Japan, Cat#R045), using pSNAP-tag(T7)-2 vector (NEB, N9181S) as a template. Then PCR product added with XhoI sites at the both ends was inserted into XhoI site of the plasmid pSN672 by using In-Fusion® HD Cloning Kit (Takara Bio Inc, Cat#639649). This KIF1A (1–393)-LZ-SNAP-6His construct was then used as template to generate the 3 aberrant motors by site-directed mutagenesis using the KOD Plus mutagenesis kit (TOYOBO, Cat#SMK-101). A recombinant human KIF1A(1–393)-LZ-SNAP-Strep-tagII construct was used to develop heterodimers of wild type and mutant KIF1A. To create this KIF1A motor domain construct, a human KIF1A(1–393)-LZ-mScarlet-I-Strep-tagII construct namely pSN643 was modified as follows: The mScarlet-I sequence of pSN643 was replaced by SNAP cDNA sequence. All constructs were confirmed by sequencing. Inactive mutant kinesin 5 B (G234A) expression vector was a gift from Dr. Yuta Shimamoto (National Institute of Genetics), which has been described (pYS05 K560 pET-17b KIF5B (1-560, G234A)::LZ::His-tag)^27^.

### Recombinant Protein Expression and purification (Homodimers)

Proteins were expressed in BL21(DE3) and SNAP-tagged KIF1A motor domain homodimers and inactive mutant KIF5B (G234A) homodimer were purified as previously described^8^. Briefly, His-tagged SNAP-tagged KIF1A motor domain plasmids were transformed into BL21(DE3) (Novagen #69450), and the cells were cultured on LB agar supplemented with ampicillin at 37°C overnight. Colonies were picked and cultured in 10 mL LB medium supplemented with ampicillin overnight. Next morning, the culture was transferred to 750 mL of 2.5×YT (20 g/L Tryptone, 12.5 g/L Yeast Extract, 6.5 g/L NaCl) supplemented with 10 mM phosphate buffer (pH 7.4) and 100 µg/mL ampicillin in 2 L flasks and shaken at 37°C. Two flasks were routinely prepared. When OD600 reached 0.6, the flasks were cooled in ice-cold water for 30 minutes, then isopropyl-β-D-thiogalactoside (IPTG) was added to the cooled culture to final concentration of 0.2 mM to induce expression. The culture was shaken at 18°C overnight (∼16 hours), and the bacteria expressing recombinant proteins were harvested by centrifugation as 4,000 rpm, 20 minutes, 4°C, (HITACH himac CR-21). The cell pellet was resuspended in Phosphate Buffered Saline; PBS(-) and centrifuged again (4000 rpm, 20 minutes, 4°C). Supernatant was discarded completely, then cell pellets were frozen and stored at -80°C until use. To purify recombinant protein, bacteria pellets were resuspended in protein buffer (50 mM HEPES, pH 8.0, 150 mM KCH_3_COO, 2 mM MgSO_4_, 10% glycerol), supplemented with 1 mM ATP, protease inhibitors, such as 4-(2-Aminoethyl) benzenesulfonylfluoride (AEBSF, FUJIFILM), Leupeptin (FUJIFILM, Wako #336-40413), Aprotinin from Bovine Lung (FUJIFILM, Wako #013-28311), and Pepstatin A (FUJIFILM, Wako #330-43973)), using 5 mL protein buffer per gram of wet cell paste. After gentle vortex, bacteria resuspension was with Benzonase nuclease (Novagen) and Lysozyme (Merch), followed by sonication using Branson SONIFIER 250. Lysate was obtained by centrifugation (40,000 rpm, 30 minutes, 4°C, HITACH himac CP-80α). After centrifugation, the supernatant was loaded on TALON® Metal Affinity Resin (Takara Bio Inc, Cat#635502), and incubated for 30 min at 4°C with mild shaking under platform shaker. The resin was washed twice with His-tag wash buffer (50 mM HEPES, pH 8.0, 450 mM KCH_3_COO, 2 mM MgSO_4_, 10% glycerol, 10 mM imidazole), containing 100 µM ATP. Protein was eluted with 10 mL His-tag elution buffer (50 mM HEPES, pH 8.0, 150 mM KCH_3_COO, 2 mM MgSO_4_, 10% glycerol, 500 mM imidazole) containing 100 µM ATP. Eluted fractions were collected and concentrated to ∼300 µL using Amicon Ultra centrifugal filters (Merck, Darmstadt, Germany). The affinity-purified protein was further separated by gel filtration using a Superdex™ 200 Increase 10/300 GL column (Cytiva) in protein buffer on an NGC Chromatography system (Bio-Rad Laboratories). Peak fractions were collected and aliquoted and snap frozen in liquid nitrogen. Protein concentration was assessed by Bradford method (BioRad Protein assay dye Reagent Concentrates, Cat#5000006). Each step of purification was analyzed by SDS-polyacrylamide gel electrophoresis (SDS-PAGE) (Fig. S1 in Supplementary Information). All recombinant DNA experiments conducted in accordance with Cartagena Protocol on Biosafety and all procedures were approved by Genetic Modification Safety Committee of Yamaguchi University (J18016). Preparation of all plasmids and recombinant proteins were performed at Yamaguchi University Center for Gene Research.

### Purification of heterodimers

BL21(DE3) cells transformed with KIF1A(1–393)::LZ::SNAP:: Strep-tag II plasmid were cultured in LB supplemented with kanamycin at 37°C. Competent cells were prepared using a Mix&Go kit (Zymogen). The competent cells were further transformed with mutant KIF1A(1–393)::LZ::SNAP::His plasmid and selected on LB agar supplemented with ampicillin and kanamycin. Colonies were picked and cultured in 10 mL LB medium supplemented with ampicillin and kanamycin overnight. Next morning, 10 mL of the medium was transferred to 750 mL 2.5×YT supplemented with carbenicillin and kanamycin in a 2 L flask and shaken at 37°C. Two flasks were routinely prepared. The procedures for protein expression in bacteria and preparation of bacterial lysate were the same as for the purification of homodimers except that the component of purification buffer was 50 mM HEPES, pH 8.0, 150 mM KCH_3_COO, 2 mM MgSO_4_, 1 mM EGTA, 10% glycerol. Lysate was loaded on Strep-Tactin® resin (IBA, GmbH) (bead volume: 2 ml) and incubated for 1 hour at 4°C with mild shaking under platform shaker. The resin was washed with 40 mL wash buffer. Protein was eluted with 10 mL protein buffer supplemented with 2.5 mM d-Desthibiotin (Merck, Sigma-Aldrich, D1411). Eluted solution was then loaded on Ni-NTA agarose (QIAGEN, #30210) (bead volume: 2 mL) and incubated for 1 hour at 4°C with mild shaking under platform shaker. The resin was washed with 40 mL His-tag wash buffer (50 mM HEPES, pH 8.0, 450 mM KCH_3_COO, 2 mM MgSO_4_, 10 mM imidazole, 10% glycerol), containing 100 µM ATP. Protein was eluted with His-tag elution buffer (50 mM HEPES, pH 8.0, 150 mM KCH_3_COO, 2 mM MgSO_4_, 10% glycerol, 500 mM imidazole) containing 100 µM ATP. Eluted fractions were collected and concentrated to ∼300 µL using Amicon Ultra centrifugal filters (Merck), then subjected to separation on NGC chromatography system (Bio-Rad) equipped with a Superdex™ 200 Increase 10/300 GL column (Cytiva). Peak fractions were pooled, concentrated in an Amicon filter again, and snap frozen in liquid nitrogen. The concentration and quality of the protein were assessed by Bradford method and SDS-PAGE, respectively (Fig. S2 in Supplementary Information).

### Microtubule preparation

Tubulin was purified from porcine brain (Tokyo Shibaura Organ Co., Ltd.) through two cycles of polymerization and depolymerization using the revised method of referece^41^, and stored at −80°C in PEM buffer (100 mM PIPES, 1 mM MgCl_2_, 1 mM EGTA, pH 6.9, adjusted with KOH). For polymerization, the tubulin solution was mixed with 2 mM GTP, 10 mM MgCl_2_, and 15% DMSO, and was then incubated at 37°C for 30 minutes. Prior to polymerization, the tubulin solution was centrifuged at 180,000 × g for 5 minutes at 2°C, and the supernatant was collected. The polymerized solution was then incubated at room temperature for 20 minutes in BRB80 buffer (80 mM PIPES, 1mM MgCl_2_, 1mM EGTA, pH 6.8 w/KOH) supplemented with 10 *μ*M Taxol (163-28163; FUJIFILM) to stabilize the microtubules. The microtubules were pelleted by centrifugation and then resuspended in BRB80 buffer. Microtubules were stored in the dark at room temperature for up to 2 weeks.

### Preparation of nanosprings (NSs) and DNA calibration rod

NSs were designed using caDNAno software, as described in the method from the reference^21^. Briefly, NSs were designed by introducing a negative superhelical strain into a four-helix bundle arranged on a square lattice. To impose the torsional strain, two additional nucleotides were periodically inserted every 32 bp in the upper pair of helices, while two nucleotides were deleted at the corresponding positions in the lower pair of helices. This design yielded a uniform coil structure with a diameter of ∼35 nm. For the kinesin-NS conjugation, 32-base single stranded DNA (ssDNA) handles (handle 32A and 32B in Table S2 in Supplementary Information) were extended from both ends of the NS. To construct the NS, 10 nM scaffold (p8064, tilibit nanosystems) was mixed with 100 nM core staples (Table S3), 100 nM handle staples (Table S2) and 12.4 mM Cy3-labeled antihandle (Table S4). The folding reaction was carried out in folding buffer (5 mM Tris pH 8.0, 1 mM EDTA and 14 mM MgCl_2_) with rapid heating to 80°C and cooling in single degree steps to 60°C over 2 h followed by additional cooling in single degree steps to 25°C over another 24 h. DNA calibration rods were also prepared as previously described^21^. Briefly, 10 nM scaffold (p8064) was mixed with 100 nM core staples (Table S5), 100 nM handle staples (Table S6), 12.4 mM Cy3-labeled antihandle and 800 nM biotin-labeled antihandle (Table S7). The folding reaction was carried out in folding buffer (5 mM Tris pH 8.0, 1 mM EDTA and 14 mM MgCl_2_), and the heating and cooling condition was the same as that for the NS. The folded DNA nanostructures were purified by glycerol gradient ultracentrifugation as previously described^21^. Briefly, 15%–45% (v/v) gradient glycerol solutions in buffer A (1×TE buffer containing 11 mM MgCl_2_) were made, and the glycerol fractions containing monomeric nanostructures were determined by agarose gel electrophoresis. The concentration of the nanostructures was determined with a Nanodrop spectrophotometer (Thermo Scientific), and the solution was aliquoted and stored at −80 °C until use. To minimize electrostatic interference between NSs and microtubules, NSs were coated with oligolysine conjugated to polyethylene glycol (K10 PEG5K, purchased from Alamanda Polymers)^42^ as follows. 10 microliters of 4.2 nM NSs were mixed with 0.45 µL of K10–PEG5K at a concentration corresponding to an N:P ratio (ratio of nitrogen in amines to phosphates in DNA) of 0.5:1. The mixture was then incubated at room temperature for 30 minutes.

### Kinesin-NS conjugation

A 32-base amine-modified DNA oligonucleotide (Hokkaido System Science) (oligo 32A* and 32B* in Table S2 in Supplementary Information) was conjugated with BG-GLA-NHS (S9151S; New England Biolabs) through the reaction between the ester and amine groups (BG-oligonucleotide). Here, the oligo 32A* and 32B* was complementary to the ssDNA handle sequence (handle 32A and 32B) of the NS. Then, the 35 µM BG-oligonucleotide (2 µL) and 1 µM SNAP-tagged KIF1A (50 µL) or 1 µM SNAP-tagged inert KIF5B mutant (G234A) (50 µL) were mixed and incubated at room temperature for 1 hour. The oligo-labelled kinesin was aliquoted and stored at -80°C. When the oligo-labeled kinesin and NS were conjugated, 5 µL of oligo 32A*-labeled KIF1A or oligo 32B*-labeled inert KIF5B (0.5–1 µM) was mixed with 10 µL of NSs (∼5 nM) in SRP90 buffer (90 mM HEPES, 50 mM CH_3_COOK, 2 mM Mg(CH_3_COO)_2_, 1 mM EGTA, pH 7.6, adjusted with KOH), and incubated on ice for 30 minutes.

### Single molecule experiment on kinesin-NS conjugates

A flow chamber was constructed from two uncoated coverslips of different sizes: an 18 mm × 18 mm coverslip (2918COVER18-18, Iwaki) was placed on top, and a 24 mm × 24 mm coverslip (C024241, Matsunami) was placed at the bottom, separated by two spacers of ∼50 *μ*m thickness, resulting in a volume of approximately 10 µL. Unless otherwise noted, the assay buffer was the SRP90 buffer supplemented with 20 µM Taxol, 0.4 mM 2-mercaptoethanol (131-14572, FUJIFILM), and 0.2 mg/mL casein (C5890-500G, Sigma-Aldrich). Microtubules were diluted in the buffer (without casein), and flowed into the chamber, allowing them to attach to the glass surface of the chamber through non-specific interaction during a 5-minute incubation. Then, the chamber was incubated with the blocking buffer (assay buffer containing 0.5 mg/mL casein) for 5 minutes to prevent nonspecific binding, during which microtubules that did not adhere to the glass surface were washed away using the blocking buffer. Next, the assay buffer containing kinesin-NS conjugates was flowed into the chamber, followed by a 5-minute incubation to allow the kinesin-NS conjugates to attach to the microtubules. Finally, the assay buffer, additionally supplemented with 1 mM ATP, 3 mg/mL glucose, 0.1 mg/mL glucose oxidase, and 0.02 mg/mL catalase, was flowed into the chamber. Note that the latter three regents are an oxygen scavenger system for fluorescence antifade protection. This step allowed for the removal of kinesin-NS conjugates that had not attached to the microtubules. Finally, to prevent the evaporation of the buffer inside the chamber during the experiment, the open ends of the flow chamber were sealed with clear nail polish. The motion of kinesin-NS conjugates was observed using a fluorescence microscope (IX83, EVIDENT) equipped with a 100× oil immersion objective lens (UPlanFL N 100×/1.30 Oil, EVIDENT) and a fluorescence mirror unit (TRITC-B, Semrock), and was recorded at 33 frames per second (fps) with an effective pixel size of 65 nm by a CMOS camera (ORCA-Flash4.0, Hamamatsu Photonics). Experiments were also performed at 100 fps in addition to the recording rate of 33 fps (Supplementary Information). The flow chamber was maintained at 25°C using a heating plate (HP-R-Z002, Live Cell Instrument) during observation.

### Observation of DNA calibration rod

A flow chamber was prepared in the same manner as described for the observation of kinesin-NS conjugates. Biotinylated BSA (0.5 mg/mL, 29130, Thermo Scientific) was incubated for 5 minutes to coat the glass surface of the chamber. Excess biotinylated BSA was washed by the assay buffer (SRP90 buffer supplemented with 10 mM Mg(CH_3_COO)_2_). Subsequently, neutravidin (0.5 mg/mL, 31000, Thermo Scientific) was flowed into the chamber and incubated for 3 minutes. Unbound neutravidin was removed by the assay buffer. Finally, ∼6 pM biotinylated DNA calibration rods^21^ were flowed into the chamber and incubated for 5 minutes to allow their attachment to the glass surface. Unbound DNA calibration rods were washed out by the assay buffer containing the oxygen scavenger system for fluorescence antifade protection (3 mg/mL glucose, 0.1 mg/mL glucose oxidase, and 0.02 mg/mL catalase).

### Conversion from image to force

Video acquisition was conducted using the High Speed Recording software (Hamamatsu Photonics), and the exported 8-bit video data in uncompressed AVI format were imported into a custom-written Python program for further analysis. The fluorescent spots corresponding to the DNA calibration rods and NSs were analyzed using the chain fit method (equation (2)). To estimate their lengths (*L*) using equation (2), parameter fitting was performed using curve fitting via the ‘curve_fit’ function from the ‘scipy.optimize’ module of Python (version 3.11.3). Data acquisition of the NS fluorescence images was performed under the same optical conditions (e.g., excitation laser intensity, dichroic mirror, filter, and camera settings) as those used for the DNA calibration rod. A correction was applied to the NS’s length (*L*) to account for possible bending of the DNA origami structure due to thermal fluctuations (Fig. 2e and 2f). The modified length 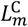 is calculated as 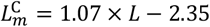 where *L* is the estimated value by the chain fitting method (equation (2)). For the parameter *σ*_long_ estimated by the Gaussian fitting method (equation (1)), the length 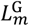 is calculated as 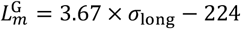. These empirical linear relationships between 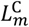 and *L*, and between 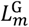 and *L* were obtained from the DNA calibration rod experiment presented in Fig. 2f. Then, the force value corresponding to the length value (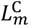 or 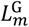) was determined from the force-extension curve (Fig.1d of the main text). Here, the force-extension curve was fitted with an exponential function (*F*(*x*) = 4.08 × 10^−4^ ∙ exp (16.6 ∙ *x*)), where *x* is the extension in *μ*m and *F*(*x*) is the force in pN.

### Angular fluctuation

Let *θ*_*t*_ denote the angle between the major axes of a microtubule and a NS images (the major axis of the ellipse fitted to the fluorescence image of the NS) in frame *t*. Then angle *θ*_*t*_ is converted to *ϕ*_*t*_ = 2*θ*_*t*_ ∈ [−*π, π*], and mapped to a unit vector 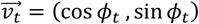. The mean of 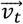 over 10 frames is defined as 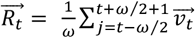. The angular fluctuation is quantified as 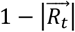. Here, the relation 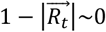 indicates that a NS is aligned with the direction of a microtubule, while 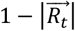 approaching 1 indicates that the NS is oriented away from the direction of the microtubule. We considered the NS to be attached to the microtubule when 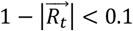. If this condition lasted for at least 5 consecutive frames, the corresponding time interval was measured as the attachment duration.

### Rate of relative increase in NS’s length

The attachment of KIF1A to the microtubule was identified by the marked suppression of angular fluctuations of the NS relative to the microtubule, which clearly distinguished the bound state from the diffusive search behavior (e.g., red regions in Fig. 3a). Within this attachment duration, the force plateau was detected by analyzing the temporal change in NS length (e.g., violet regions in Fig. 3a): the 10-frame moving average of the extension, 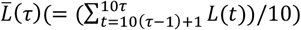 where *t* represents a video frame, was computed, and the regions where the relative increase rate 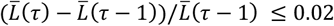 were regarded as plateau segments. The mean NS extension over these plateau segments was defined as *L*_stall_.

### Statistical analysis

Statistical analyses were performed using Python (version 3.11.3) with scipy.stats (for Shapiro–Wilk, Levene, Mann–Whitney U, and Kruskal–Wallis tests) and scikit-posthocs (for Dunn’s post hoc test with Bonferroni correction). Statistical comparisons were conducted for *L*_stall_ and Δ*t* measurements. The normality of each dataset was assessed using the Shapiro–Wilk test. All datasets were non-normally distributed, except for *L*_stall_ values obtained from the homozygous P305L mutant (P305L/P305L) and heterozygous V8M mutant (V8M/WT). Homogeneity of variances was evaluated using Levene’s test. Homogeneity of variance was confirmed for *L*_stall_ between the w/o PEG and w/PEG conditions, and among WT/WT and A255V genotypes. However, variance differed significantly among WT/WT and P305L genotypes and among WT/WT and V8M genotypes. All Δ*t* datasets showed unequal variances. Based on these results, non-parametric statistical tests were applied. The Mann–Whitney U test was used for comparing *L*_stall_ between w/o PEG and w/PEG, and for comparing Δ*t* between homozygous and heterozygous mutant combinations. The Kruskal–Wallis test was applied to compare *L*_stall_ across WT/WT, mutant/mutant, and mutant/WT combinations for each KIF1A mutant (P305L, V8M, A255V), followed by Dunn’s test with Bonferroni correction.

## Data availability

The data that support the findings of this study are available from the corresponding author upon reasonable request.

## Acknowledgements

We acknowledge Dr. Yuta Shimamoto for providing plasmids, and Ms. Kimiko Nagino and Mr. Gai Ohashi for purifying microtubules. This work was supported by JSPS KAKENHI (Grant No. 23H02442), the Precise Measurement Technology Promotion Foundation (PMTP-F), and a KIF1A.org Mini Grant to K.H, JSPS KAKENHI (Grant No. 21H01053) and JST, CREST (Grant No. JPMJCR2023) to M.I., JST PRESTO (Grant No. JPMJPR21E2) to T.A.

## Author contributions

N.T. performed the single-molecule experiments and data analysis. H.F. and T.A. contributed to protein purification. M.I. provided DNA origami materials and associated protocols. K.H. designed the experiments together with M.I. and T.A., and wrote the manuscript in collaboration with M.I. and N.T.

