## Supplementary text for "Stall force measurement of the kinesin-3 motor KIF1A using a programmable DNA origami nanospring"

(A) KIF1A (WT) homodimer

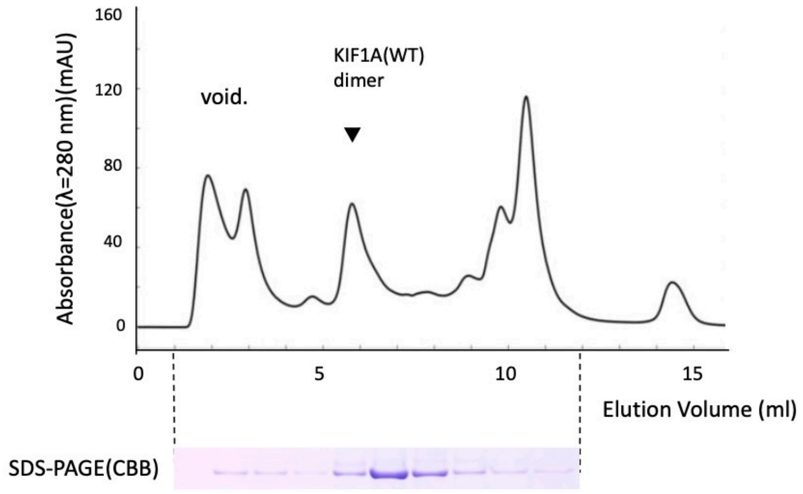


KIF1A (V8M) homodimer

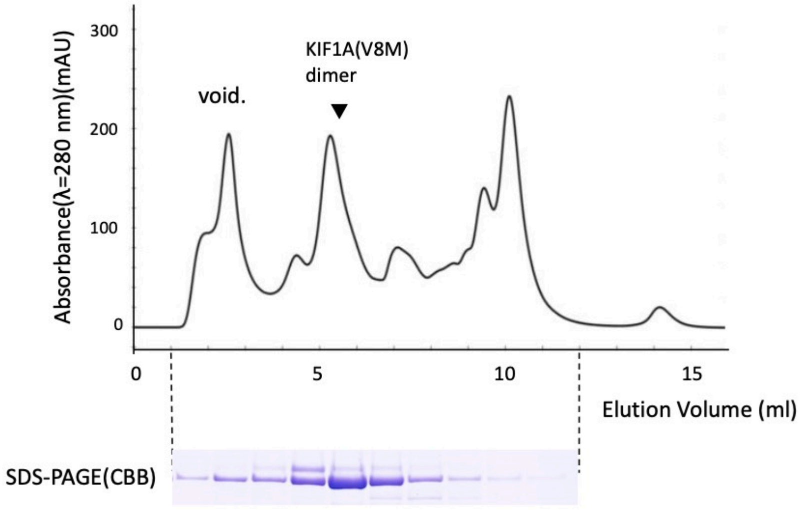


KIF1A (A255V) homodimer

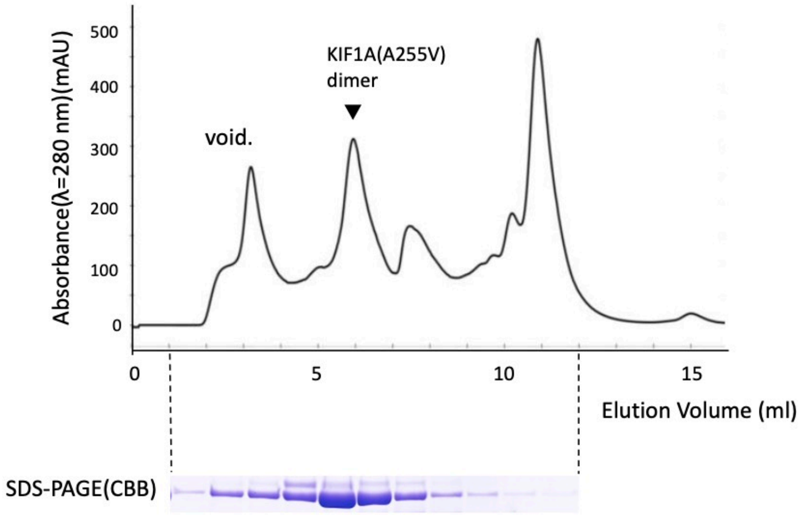

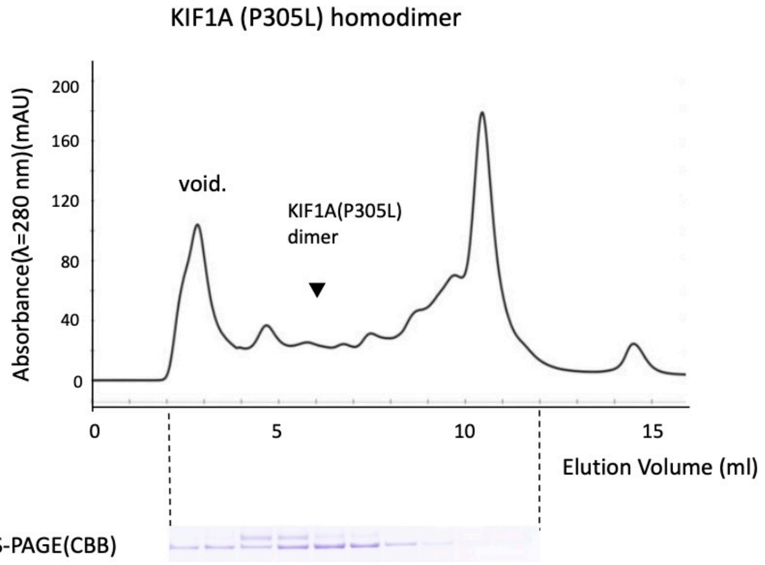


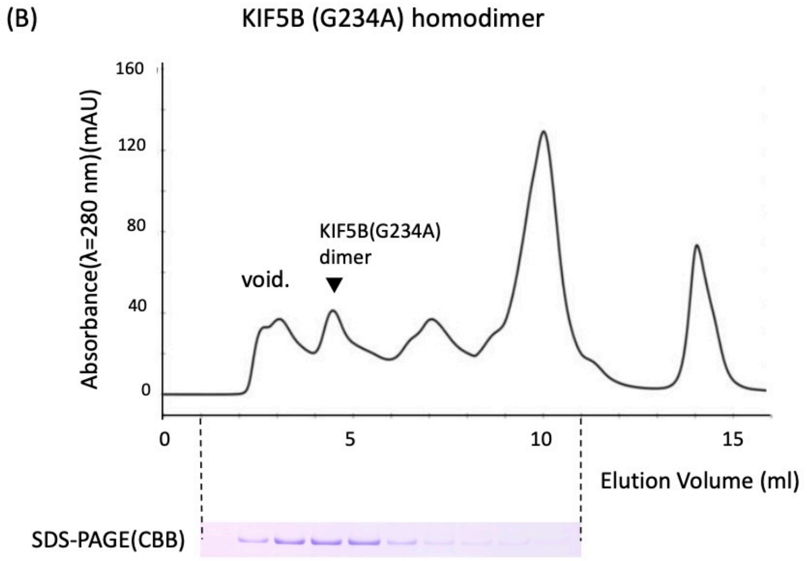


**Fig. S1 Purification of recombinant SNAP tagged KIF1A and KIF5B proteins.** (A) Size-Exclusion Chromatogram for SNAP tagged wild type or mutant KIF1A dimerized motors (1-393-LZ). Elution positions of prep contaminants (void.) are noted. Dimerized KIF1A motor domain was shown by arrow heads. Below: Coomassie-stained SDS-polyacrylamide gel electrophoresis (PAGE) of the column elution fractions. (B) Size Exclusion SDS-PAGE NGC Chromatogram showing elution profiles of KIF5B (G234A).

KIF1A (WT/V8M) heterodimer

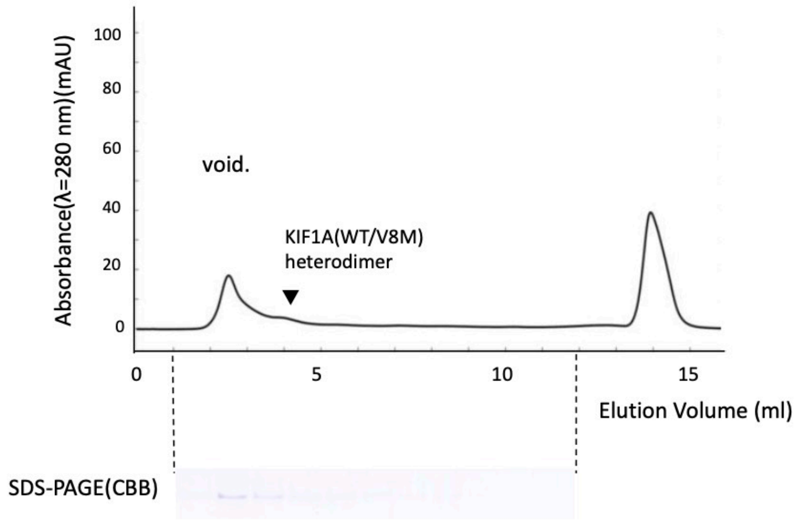


KIF1A (WT/A255V) heterodimer

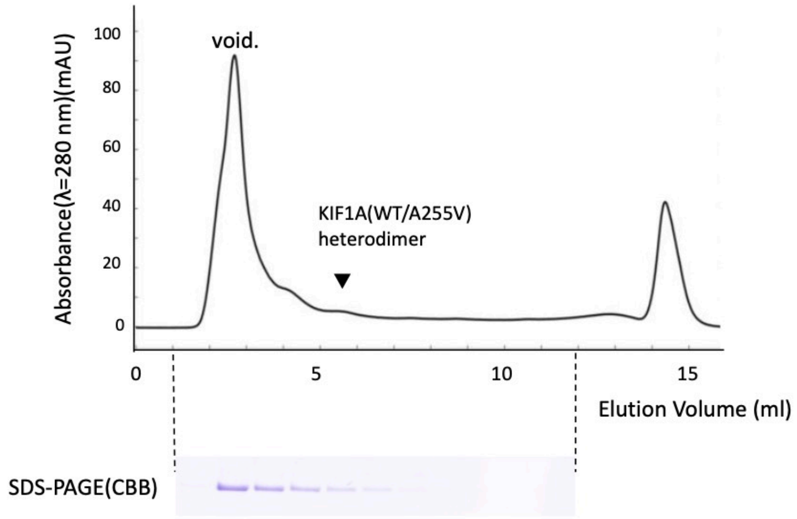


KIF1A (WT/P305L) heterodimer

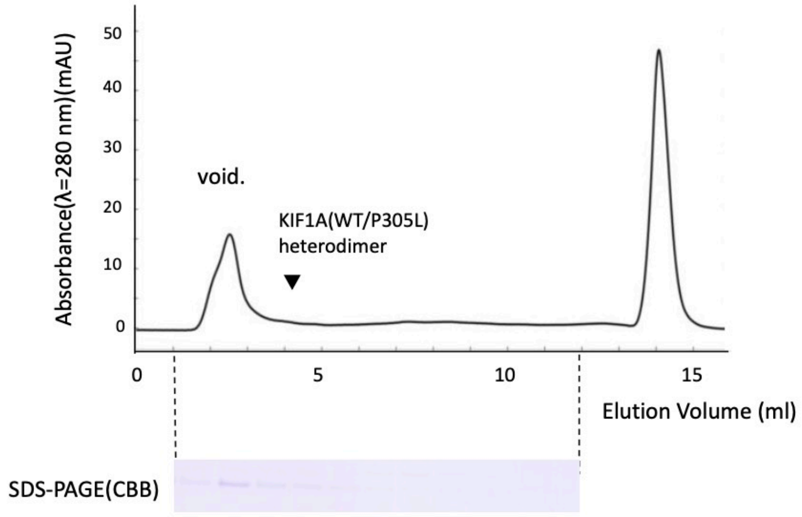

**Fig. S2 Purification of recombinant SNAP tagged KIF1A heterodimers.** Results of Size Exclusion Chromatogram for heterodimers composed of wild type and mutant KIF1A. Elution positions of prep contaminants (void.) and heterodimerized KIF1A motor domain are noted. Below: Coomassie-stained SDS-PAGE of the column elution fractions showing dimerized KIF1A.

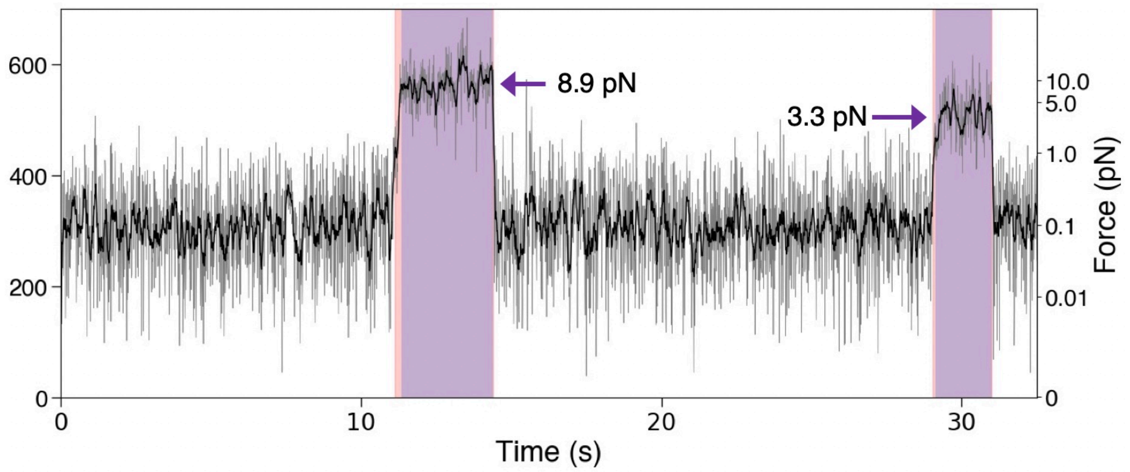


**Fig. S3** Time course of a nanospring (NS) extension at a recording rate of 100 fps for wild-type KIF1A homodimers. No significant difference in $L_{\mathrm{stall}}$ was observed between recordings at 33 fps (Fig. 3, main text) and 100 fps. The lighter-colored regions in the graph represent the attachment durations, while the darker-colored regions indicate the stall durations. The identification of these durations is described in the Methods section.

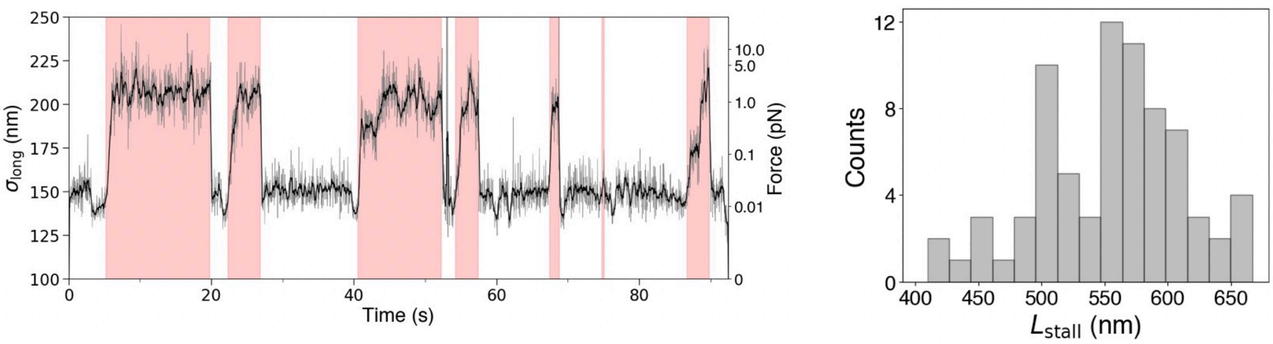


**Fig. S4 Estimation of NS extensions using the Gaussian fitting method** (equation (1), main text). The conversion from $\sigma_{\mathrm{long}}$ to $L_m^{\mathrm{G}}$ was performed using the relation $L_m^{\mathrm{G}} = 3.67 \times \sigma_{\mathrm{long}} - 224$ (Methods, main text). The $L_{\mathrm{stall}}$ estimated by the Gaussian fitting method was 534 ± 58 (SD) nm.

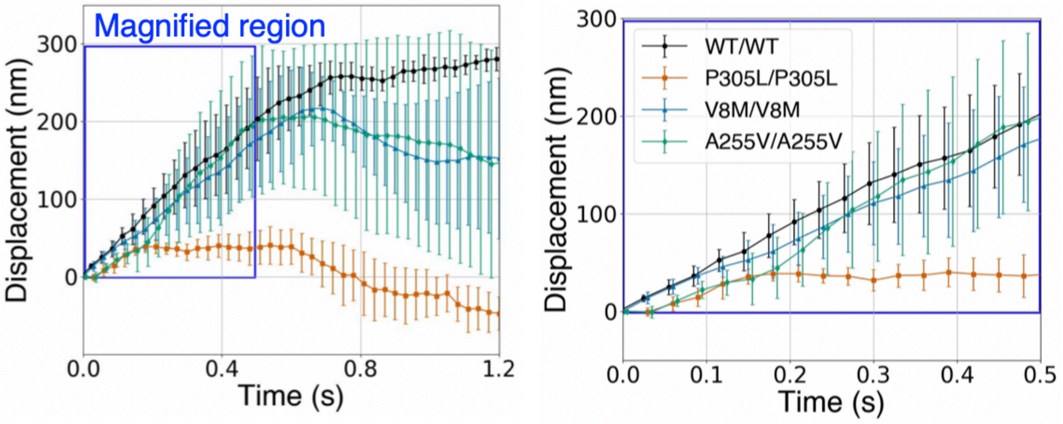


**Fig. S5 Displacement (calculated from the extensions of NSs) for KIF1A KAND mutants.** Approximately 10 stall events were overlaid. V8M and A255V are slightly slower than WT, whereas P305L is significantly slower.

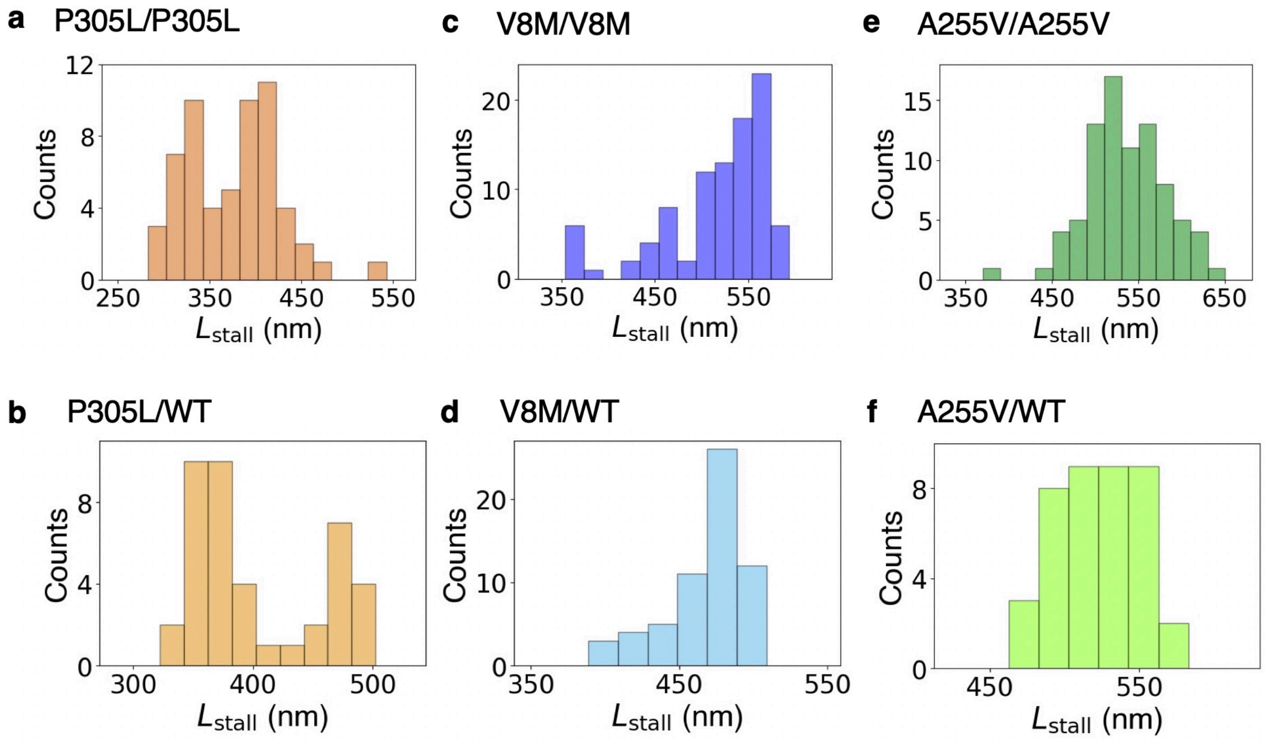


**Fig. S6 Histograms of $L_{stall}$ for KIF1A KAND mutants.** The time courses of the NS extensions are presented in Fig. 4 of the main text. The mean values of $L_{stall}$ are listed in Table 1 of the main text. Stall events: $n$=58 for P305L/P305L, $n$=41 for P305L/WT, $n$=95 for V8M/V8M, $n$=61 for V8M/WT, $n$=83 for A255V/A255V, and $n$=40 for A255V/WT.

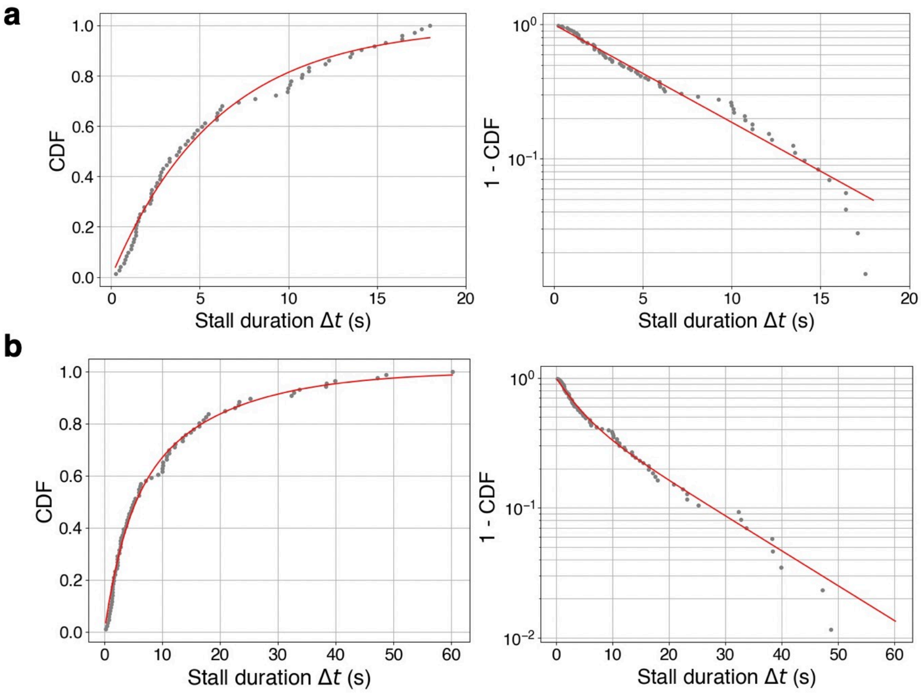


**Fig. S7 Cumulative distributions of stall duration ($\Delta t$) in the case of the KIF1A(WT).** (**a**) When $\Delta t \leq 20$ s, the data were fitted using $1 - e^{-kt}$ with $k = 0.17$. (**b**) When fitting over a wide range of $\Delta t$, the data were fitted using $Ak_1e^{-k_1t} + (1-A)k_2e^{-k_2t}$ with $k_1 = 0.26$, $k_2 = 0.062$ and $A = 0.44$.

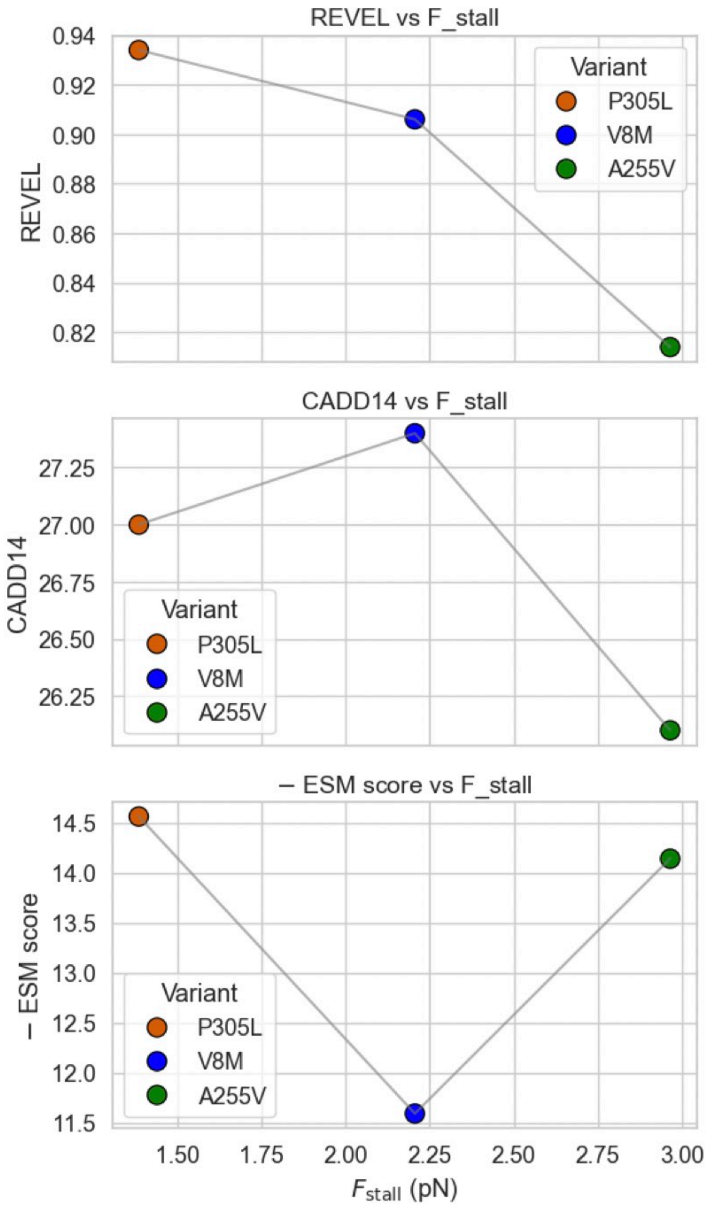


| | P305L | V8M | A255V |
|---|---|---|---|
| REVEL | 0.934 | 0.906 | 0.814 |
| CADD14 | 27.0 | 27.4 | 26.1 |
| ESM score | -14.562 | -11.59 | -14.14 |

**Fig. S8 Comparison between stall force and KAND severity scores.** These scores are cited from the previous study [Boyle *et al*., *HGG Advance* **2**, 100026 (2021)].

**Table S1**

**Plasmid list**

| Plasmid | Description | Backbone |
|---|---|---|
| pSN643 | KIF1A(1-393)::LZ::mScarlet-I::Strep-tag II | pET28a |
| pSN672 | KIF1A(1-393)::LZ::His-tag | pET21a |
| pHF001 | KIF1A(1-393)::LZ::SNAP::His-tag | pET21a |
| pHF002 | KIF1A(1-393)(P305L)::LZ:: SNAP::His-tag | pET21a |
| pHF003 | KIF1A(1-393)(V8M)::LZ:: SNAP::His-tag | pET21a |
| pHF004 | KIF1A(1-393)(A255V)::LZ:: SNAP::His-tag | pET21a |
| pHF005 | KIF1A(1-393)::LZ::SNAP:: Strep-tag II | pET28a |
| pYS05 | KIF5B(1-393)(G234A)::RA::SNAP::His-tag | pET17b |

RA: RA linker

**Table S2**

**Handle staples and anti handles used to link KIF1A and inert KIF5B at the end of NSs.**

| **Name** | **Sequence (5' to 3')** |
|---|---|
| handle 32A staple | AATCGGAACCCTAAAGGAAAAACCGTCTATCA<br>*CCCACCTATTTTCACCCCACCCTTCCCCCAAC* |
| handle 32B staple | *CCTTATCCCAAACCCCTCCCCCAACCATTCCC*<br>TTACCAGTATAAATGAGTAATGTGTAGGTAAA |
| oligo 32A* | $NH_2$/GTTGGGGGAAGGGTGGGGTGAAAATAGGTGGG |
| oligo 32B* | GGGAATGGTTGGGGGAGGGGTTTGGGATAAGG/$NH_2$ |

In the sequences of staples handle 32A and 32B, italicized regions indicate single-stranded DNA (ssDNA) handle sequences.

**Table S3**

Core staples and handle staples for Cy3 to build the NS. Sequences in italics indicate the handle site.

| **Sequence (5' to 3')** | **Name** |
|---|---|
| ATAAGTGCCGTGGGGAAAGCTATTAAAGAACGTGGAC | Core Staple_01 |
| ATTAGCGGGGTAAGGGAAGGTTGAGTGTTGTTCCAGT | Core Staple_02 |
| AGGCTGAGACTGGGCGCTGTCCCTTATAAATCAAAAG | Core Staple_03 |
| CCTATTATTCTGCGTAACCATCCTGTTTGATGGTGGT | Core Staple_04 |
| AACAGTTAATGCGCCGCTAAGCGGTCCACGCTGGTTT | Core Staple_05 |
| GTCAGTGCCTTTTGACGAGCCCTTCACCGCCTGGCCC | Core Staple_06 |
| AGGAGTGTACTTTAGAATCTCTTTTCACCAGTGAGAC | Core Staple_07 |
| GTAAGCGTCATCCGATTAAAGGCGGTTTGCGTATTGG | Core Staple_08 |
| GAAAGCGCAGTACGCCAGAAGCTGCATTAATGAATCG | Core Staple_09 |
| AACAAATAAATTCAGTGAGCTGCCCGCTTTCCAGTCG | Core Staple_10 |
| CAGGTCAGACGCCATCACGAGTGAGCTAACTCACATT | Core Staple_11 |
| GCCGCCGCCAGCTTCTTTGGAAGCATAAAGTGTAAAG | Core Staple_12 |
| CTCAGAGCCGCTGAGTAGATATCCGCTCACAATTCCA | Core Staple_13 |
| CTCAGAACCGCCTGGTAATATCATGGTCATAGCTGTT | Core Staple_14 |
| ACCGGAACCGCGCCATTGCTTGAGGATCCCCGGGTAC | Core Staple_15 |
| ATAATCAAAATATACCTACTGCCTGTTCTTCGCGTCC | Core Staple_16 |
| AGCCCCCTTATAATGGATTTCACGGTCATACCGGGGG | Core Staple_17 |
| TAGCGCGTTTTGTCACACGTGTGGTGCTGCGGCCAGA | Core Staple_18 |
| AGAATCAAGTTTGGCCAACATCCAGCGCAGTGTCACT | Core Staple_19 |
| ATCGATAGCAGGAAAGCGTCAGCCAGCGGTGCCGGTG | Core Staple_20 |
| AGCAAGGCCGGATTTTTGAAAATCGTTAACGGCATCA | Core Staple_21 |
| AGCAAAATCACCGAACTGACGTCATAAACATCCCTTA | Core Staple_22 |
| ACCGACTTGAGAAAATACCCCTGCGGCTGGTAATGGG | Core Staple_23 |
| ATTATTCATTAAAAACAGAGGCGCGGTTGCGGTATGA | Core Staple_24 |
| TTGAGGGAGGGCCGCCTGCGGTCATTGCAGGCGCTTT | Core Staple_25 |
| GCCAAAGACAAGCAGCAAACCAGCTTACGGCTGGAGG | Core Staple_26 |
| ATCAATAGAAATGCTGAACACGGCAGCACCGTCGGTG | Core Staple_27 |
| GACACCACGGAAATATCTGGCTGGTCTGGTCAGCAGC | Core Staple_28 |
| AAAGGTGGCAAGAAAGGAAGGAACGTGCCGGACTTGT | Core Staple_29 |
| TATGTTAGCAATCTTTAGGCTGGCAGCCTCCGGCCAG | Core Staple_30 |
| CTGGCATGATTGAGCCGTCTCCGGCAAACGCGGTCCG | Core Staple_31 |

| | |
|---|---|
| AACGCAATAATTTAGAAGTATGAAGGGTAAAGTTAAA | Core Staple_32 |
| CGAACAAAGTTACAACTCGCCCGTAAAAAAAGCCGCA | Core Staple_33 |
| AGCCCTTTTTATTATTAATAGTTGGGCGGTTGTGTAC | Core Staple_34 |
| ACAATGAAATATCATTTTGATCAAACTTAAATTTCTG | Core Staple_35 |
| TTGAGTTAAGCGGAGCGGAAGCTCTCACGGAAAAAGA | Core Staple_36 |
| CGCTAATATCATCAGATGATGTGAGAGATAGACTTTC | Core Staple_37 |
| CACCCTGAACATGATTGTTAACGTACAGCGCCATGTT | Core Staple_38 |
| GAAGCGCATTAGAAGGGTTGGGAACGGATAACCTCAC | Core Staple_39 |
| CCTTTACAGAGTTTGCACGGCCAGTGCCAAGCTTTCA | Core Staple_40 |
| TTTGTTTAACGCGTAGATTAGGGTTTTCCCAGTCACG | Core Staple_41 |
| ATTTATCCCAATATACAGTGATGTGCTGCAAGGCGAT | Core Staple_42 |
| TTTGCCAGTTAGAAACAATGCCTCTTCGCTATTACGC | Core Staple_43 |
| AACGCTAACGAGAATACCAGCTGCGCAACTGTTGGGA | Core Staple_44 |
| CCCAGCTACAAGAATTATTCCGGAAACCAGGCAAAGC | Core Staple_45 |
| CCTTAAATCAAGAAGATGACACTCCAGCCAGCTTTCC | Core Staple_46 |
| GAACCTCCCGAAAATTAATGGGGACGACGACAGTATC | Core Staple_47 |
| CGGTATTCTAAATTACCTTTGGGCGCATCGTAACCGT | Core Staple_48 |
| AGCAAGCAAATATCAATATTTGACCGTAATGGGATAG | Core Staple_49 |
| TTCATCGTAGGCTGTAAATACCCGTCGGATTCTCCGT | Core Staple_50 |
| CTCATCGAGAACTTAGAATAGCTTTCATCAACATTAA | Core Staple_51 |
| CCAAGAACGGGTAGATTAATCAAAAATAATTCGCGTC | Core Staple_52 |
| ATCAATAATCGGAATTTATAAATCAGCTCATTTTTTA | Core Staple_53 |
| CCATCCTAATTTACCTTTTAATATTTTGTTAAAATTC | Core Staple_54 |
| AGATAAGTCCTTATATAACGATTGTATAAGCAAATAT | Core Staple_55 |
| TAATGCAGAACTCCAATCGCGGTTGATAATCAGAAAA | Core Staple_56 |
| CAGACGACGACCTTTTTCAAATCGTAAAACTAGCATG | Core Staple_57 |
| AGTACCGACAACTTCTGACGAGTCTGGAGCAAACAAG | Core Staple_58 |
| CATTTTCGAGCCCGACCGTAGAGATCTACAAAGGCTA | Core Staple_59 |
| AACAACGCCAAAGAATAAAGATAAATTAATGCCGGAG | Core Staple_60 |
| ACAGTAGGGCTTAAAAAGCCTATCACCATCAATATGATA | Core Staple_61 |
| TTTAGAGCTTGACCGAGAGGGCCGCCACCCTCAGAACCG | Core Staple_62 |
| GGCGAGAAAGGTTTGCTCACACCCTCAGAGCCACCAC | Core Staple_63 |
| AGCGGGCGCTACCTCAAGAAGCAAGCCCAATAGGAAC | Core Staple_64 |

| | |
|---|---|
| GGTCACGCTGCGAAACATGTGAGTTTCGTCACCAGTA | Core Staple_65 |
| CGCGCTTAATGCCCCCTGCTAGCATTCCACAGACAGC | Core Staple_66 |
| CTATGGTTGCTGAGTAACAACGATCTAAAGTTTTGTC | Core Staple_67 |
| TGCTTTCCTCGGGTAATAAGTAAATGAATTTTCTGTA | Core Staple_68 |
| TAAACAGGAGGACATGGCTAACTTTCAACAGTTTCAG | Core Staple_69 |
| ACAGGAACGGTCTCTGAATAGGAACAACTAAAGGAAT | Core Staple_70 |
| TGTTTTTATAACCTCATTATTCACGTTGAAAATCTCC | Core Staple_71 |
| AAAGAGTCTGTATTGGCCTAAGGAGCCTTTAATTGTA | Core Staple_72 |
| TTGTAGCAATACATTGACACTTTCGAGGTGAATTTCT | Core Staple_73 |
| CATCACTTGCCCACCAGAAGATAGTTGCGCCGACAAT | Core Staple_74 |
| TATCGGCCTTGCACCCTCACACGCATAACCGATATAT | Core Staple_75 |
| TATTACCGCCACTCCCTCAGCAGGGAGTTAAAGGCCG | Core Staple_76 |
| CGCTCATGGAACACCGGAAACCCTCAGCAGCGAAAGA | Core Staple_77 |
| CAATCGTCTGATAGCGTTTGTAGCAACGGCTACAGAG | Core Staple_78 |
| CAGATTCACCACATCGGCAACTTTTTCATGAGGAAGT | Core Staple_79 |
| AAGGGACATTCTGCCTTTAAAATACGTAATGCCACTA | Core Staple_80 |
| CCTTCTGACCTCACCGTAAAAACGAAAGAGGCAAAAG | Core Staple_81 |
| GCACAGACAATAAACGTCACATCTTTGACCCCCAGCG | Core Staple_82 |
| TCTTTAATGCGCAGTAGCAAACAAAGTACAACGGAGA | Core Staple_83 |
| CATCGCCATTACCATTTGGATAAATTGTGTCGAAATC | Core Staple_84 |
| CAGCAGAAGATAAGGTGAATTACTTAGCCGGAACGAG | Core Staple_85 |
| CAGTATTAACAAAGGTAAAATAAGGGAACCGAACTGA | Core Staple_86 |
| GCTGAGAGCCAAAGGGCGAACAGATGAACGGTGTACA | Core Staple_87 |
| AAGCATCACCTATTCATATTGGCTGACCTTCATCAAG | Core Staple_88 |
| AACCCTCAATCATAAGTTTACCGGATATTCATTACCC | Core Staple_89 |
| AATCAACAGTTCATATAAAGCTGCTCATTCAGTGAAT | Core Staple_90 |
| TATCTAAAATAACGTAGAAAGAAACACCAGAACGAGT | Core Staple_91 |
| CTAATAGATTAAAGACTCCGATGGTTTAATTTCAACT | Core Staple_92 |
| CATTTGAGGATAACGGAATACCTTATGCGATTTTAAG | Core Staple_93 |
| CAAACAATTCGACCAGAAGCAGTCAGGACGTTGGGAA | Core Staple_94 |
| TTGCCCGAACGAGAAAAGTTAAAACGAACTAACGGAA | Core Staple_95 |
| GAGTAACATTAGCAATAGCAGAAAGATTCATCAGTTG | Core Staple_96 |
| AACCACCAGAACCAATAATCATTCAACTAATGCAGAT | Core Staple_97 |

| | |
|---|---|
| ATTCCTGATTAGAGAGATAAATTACGAGGCATAGTAA | Core Staple_98 |
| CAATATAATCCAAGTCAGAACCCTCGTTTACCAGACG | Core Staple_99 |
| TCTGAATAATGGACGGGAGTAGCGAGAGGCTTTTGCA | Core Staple_100 |
| TATCAAAATTAAGAATAACAGGGGGTAATAGTAAAAT | Core Staple_101 |
| TAAAGAAATTGTCAAAAATGTCCAATACTGCGGAATC | Core Staple_102 |
| CGTCAGATGAATCCAAATAGAATCCCCCTCAAATGCT | Core Staple_103 |
| TTACATCGGGACAAAATAAACGAGAATGACCATAAAT | Core Staple_104 |
| CTGATTGCTTTGCGTCTTTACCCTGACTATTATAGTC | Core Staple_105 |
| CGCGCAGAGGCTTTTATCCGCATCAAAAAGATTAAGA | Core Staple_106 |
| CCTGAGCAAAAGATTAGTTTCAAATATCGCGTTTTAA | Core Staple_107 |
| TCAAGAAAACACTTGCGGGAACCAGACCGGAAGCAAA | Core Staple_108 |
| ATTTCATTTGAGAACGCGATTAGAGAGTACCTTTAAT | Core Staple_109 |
| ACAGTACATAACAGATATAAGGTCATTTTTGCGGATG | Core Staple_110 |
| TAACCTTGCTTAATCATTACTGAATATAATGCTGTAG | Core Staple_111 |
| ATTAATTTTCCCAAGCAAGTATGCAACTAAAGTACGG | Core Staple_112 |
| TAGCGATAGCTTATTAAACTCCATATAACAGTTGATT | Core Staple_113 |
| GAGTCAATAGTGCTGTCTTAGTAGATTTAGTTTGACC | Core Staple_114 |
| GTCTGAGAGACTACGAGCAAAATGGTCAATAACCTGT | Core Staple_115 |
| TTAGGTTGGGTGAACAAGATTGGGGCGCGAGCTGAAA | Core Staple_116 |
| GCTGATGCAAAGCGCCTGTACTAATAGTAGTAGCATT | Core Staple_117 |
| ACGCGAGAAAAAATAAACATACAGGCAAGGCAAAGAA | Core Staple_118 |
| GTTAATTTCATAAGGTAAAATAAAGCCTCAGAGCATA | Core Staple_119 |
| GGTTTGAAATACAGTAATAACCAAAAACATTATGACC | Core Staple_120 |
| GGCGTTAAATACATGTAATGGAGAAGCCTTTATTTCA | Core Staple_121 |
| TAATTACTAGAATTGAGAATTTTAGAACCCTCATATA | Core Staple_122 |
| *CTCCTATCTCCAATCACTCCT*TCCAACGTGTTTAGTATTGATATAAGT | Handle staple for Cy3_1 |
| *CTCCTATCTCCAATCACTCCT*TTGGAACAAGAACCGCGTACCAGGCGG | Handle staple for Cy3_2 |
| *CTCCTATCTCCAATCACTCCT*AATAGCCCTCAGGGATGAAGGATTAGG | Handle staple for Cy3_3 |
| *CTCCTATCTCCAATCACTCCT*TCCGAAATCGTAACACAAAGTATTAAG | Handle staple for Cy3_4 |
| *CTCCTATCTCCAATCACTCCT*GCCCCAGCAACGCCTGCTATTTCGGAA | Handle staple for Cy3_5 |
| *CTCCTATCTCCAATCACTCCT*TGAGAGAGTTAGCGTAGTGCCCGTATA | Handle staple for Cy3_6 |
| *CTCCTATCTCCAATCACTCCT*GGGCAACAAGACGTTAGTTTTAACGGG | Handle staple for Cy3_7 |
| *CTCCTATCTCCAATCACTCCT*GCGCCAGGTGCTAAACTTTGATGATAC | Handle staple for Cy3_8 |

| | |
|---|---|
| *CTCCTATCTCCAATCACTCCT*GCCAACGCGAATAGAATTACCGTTCCA | Handle staple for Cy3_9 |
| *CTCCTATCTCCAATCACTCCT*GGAAACCTATAATTTTAAGCCAGAATG | Handle staple for Cy3_10 |
| *CTCCTATCTCCAATCACTCCT*AATTGCGTGGCTCCAATGATATTCACA | Handle staple for Cy3_11 |
| *CTCCTATCTCCAATCACTCCT*CCTGGGGTTCAGCTTGGGAGGTTGAGG | Handle staple for Cy3_12 |
| *CTCCTATCTCCAATCACTCCT*CACAACATTTGATACCCCACCACCAGA | Handle staple for Cy3_13 |
| *CTCCTATCTCCAATCACTCCT*TCCTGTGTCCATCGCCGAGCCACCACC | Handle staple for Cy3_14 |
| *CTCCTATCTCCAATCACTCCT*CGAGCTCGTGAGGCTTGAGCCGCCACC | Handle staple for Cy3_15 |
| *CTCCTATCTCCAATCACTCCT*GTGAGCCTGGATCGTCCCAGAGCCACC | Handle staple for Cy3_16 |
| *CTCCTATCTCCAATCACTCCT*TTTCTGCCGAACGAGGGCCATCTTTTC | Handle staple for Cy3_17 |
| *CTCCTATCTCCAATCACTCCT*ATGCGGCGGACTAAAGTTTTCGGTCAT | Handle staple for Cy3_18 |
| *CTCCTATCTCCAATCACTCCT*GCGCGCCTAACGGGTAGCGTCAGACTG | Handle staple for Cy3_19 |
| *CTCCTATCTCCAATCACTCCT*CCCCCTGCCCAACCTATCAGTAGCGAC | Handle staple for Cy3_20 |
| *CTCCTATCTCCAATCACTCCT*GATGCCGGAAAACACTCCAATGAAACC | Handle staple for Cy3_21 |
| *CTCCTATCTCCAATCACTCCT*CACTGGTGAAGCGCGACCATTACCATT | Handle staple for Cy3_22 |
| *CTCCTATCTCCAATCACTCCT*TAAAGGTTATCGCCTGGAATTAGAGCC | Handle staple for Cy3_23 |
| *CTCCTATCTCCAATCACTCCT*GCCGGGTCGCTCCATGTTATCACCGTC | Handle staple for Cy3_24 |
| *CTCCTATCTCCAATCACTCCT*CGCACTCAGGTCAATCTATTGACGGAA | Handle staple for Cy3_25 |
| *CTCCTATCTCCAATCACTCCT*TGTCCAGCGAAAGAGGCATTCAACCGA | Handle staple for Cy3_26 |
| *CTCCTATCTCCAATCACTCCT*GTGCCATCGCATAGGCGGTTTACCAGC | Handle staple for Cy3_27 |
| *CTCCTATCTCCAATCACTCCT*AACCGCAATGACAAGAATTTTGTCACA | Handle staple for Cy3_28 |
| *CTCCTATCTCCAATCACTCCT*AGAACGTCGTAACAAAAGAAACGCAAA | Handle staple for Cy3_29 |
| *CTCCTATCTCCAATCACTCCT*AGCACATCCCCTGACGAATACATACAT | Handle staple for Cy3_30 |
| *CTCCTATCTCCAATCACTCCT*TTTTTTCGGGGCTTGATTATTACGCAG | Handle staple for Cy3_31 |
| *CTCCTATCTCCAATCACTCCT*CGATGCTGTGTGAATTACCCAAAAGAA | Handle staple for Cy3_32 |
| *CTCCTATCTCCAATCACTCCT*CAGGCGGCCATTATACGAAACCGAGGA | Handle staple for Cy3_33 |
| *CTCCTATCTCCAATCACTCCT*ATCGACATTACGTTAAAAGCAGATAGC | Handle staple for Cy3_34 |
| *CTCCTATCTCCAATCACTCCT*CTCATTTGTTACAGGTTATCTTACCGA | Handle staple for Cy3_35 |
| *CTCCTATCTCCAATCACTCCT*GACGCAGAGAATACCAAAGAGCAAGAA | Handle staple for Cy3_36 |
| *CTCCTATCTCCAATCACTCCT*TCCGTGGTCCAAAAGGACCCACAAGAA | Handle staple for Cy3_37 |
| *CTCCTATCTCCAATCACTCCT*TACCAGTCCTATCATAGGGTAATTGAG | Handle staple for Cy3_38 |
| *CTCCTATCTCCAATCACTCCT*CGGAAACAAACCAAAAAATTAACTGAA | Handle staple for Cy3_39 |
| *CTCCTATCTCCAATCACTCCT*GAGGTGGATTTGCCAGATAAAAACAGG | Handle staple for Cy3_40 |
| *CTCCTATCTCCAATCACTCCT*ACGTTGTATGGATAGCGAAAATAGCAG | Handle staple for Cy3_41 |

| | |
|---|---|
| *CTCCTATCTCCAATCACTCCT*TAAGTTGGTATTCATTAGAAACGATTT | Handle staple for Cy3_42 |
| *CTCCTATCTCCAATCACTCCT*CAGCTGGCTTCAGAAAACAGCCATATT | Handle staple for Cy3_43 |
| *CTCCTATCTCCAATCACTCCT*AGGGCGATAGGTCTTTCCAGAGCCTAA | Handle staple for Cy3_44 |
| *CTCCTATCTCCAATCACTCCT*GCCATTCGAGCGGATTTGAATCTTACC | Handle staple for Cy3_45 |
| *CTCCTATCTCCAATCACTCCT*GGCACCGCGAAAGACTGCTATTTTGCA | Handle staple for Cy3_46 |
| *CTCCTATCTCCAATCACTCCT*GGCCTCAGTCAAAGCGAGGTTTTGAAG | Handle staple for Cy3_47 |
| *CTCCTATCTCCAATCACTCCT*GCATCTGCGGTCAGGAGGCGTTTTAGC | Handle staple for Cy3_48 |
| *CTCCTATCTCCAATCACTCCT*GTCACGTTTTGATAAGGAAGGCTTATC | Handle staple for Cy3_49 |
| *CTCCTATCTCCAATCACTCCT*GGGAACAACTTAATTGCCGCGCCCAAT | Handle staple for Cy3_50 |
| *CTCCTATCTCCAATCACTCCT*ATGTGAGCGTTTTAAACCGTTTTTATT | Handle staple for Cy3_51 |
| *CTCCTATCTCCAATCACTCCT*TGGCCTTCAGTTTCATCAAGTACCGCA | Handle staple for Cy3_52 |
| *CTCCTATCTCCAATCACTCCT*ACCAATAGTGCGAACGTCCTTATCATT | Handle staple for Cy3_53 |
| *CTCCTATCTCCAATCACTCCT*GCATTAAACATTTCGCTGTAGAAACCA | Handle staple for Cy3_54 |
| *CTCCTATCTCCAATCACTCCT*TTAAATTGATTTTCATAAAATAATATC | Handle staple for Cy3_55 |
| *CTCCTATCTCCAATCACTCCT*GCCCCAAATCAATTCTTTATCAACAAT | Handle staple for Cy3_56 |
| *CTCCTATCTCCAATCACTCCT*TCAATCATATAAATCAACATGTTCAGC | Handle staple for Cy3_57 |
| *CTCCTATCTCCAATCACTCCT*AGAATCGAATTAAGCAGTAATTCTGTC | Handle staple for Cy3_58 |
| *CTCCTATCTCCAATCACTCCT*TCAGGTCATCGGTTGTAGAGAATATAA | Handle staple for Cy3_59 |
| *CTCCTATCTCCAATCACTCCT*AGGGTAGCCTTTTGCGTTAGGCAGAGG | Handle staple for Cy3_60 |
| *CTCCTATCTCCAATCACTCCT*TTCAACCGATAAAAATTCGCCATATTT | Handle staple for Cy3_61 |
| *CTCCTATCTCCAATCACTCCT*GGCCGGAGGCAATGCCGCCAACGCTCA | Handle staple for Cy3_62 |
| *CTCCTATCTCCAATCACTCCT*CTCAGGAGCAAAGGGCGGAGCCCCCGA | Handle staple for Cy3_63 |
| *CTCCTATCTCCAATCACTCCT*CCACCCTCAGAGTCCACCGGCGAACGT | Handle staple for Cy3_64 |
| *CTCCTATCTCCAATCACTCCT*CCTCATTTGAGATAGGAAAGCGAAAGG | Handle staple for Cy3_65 |
| *CTCCTATCTCCAATCACTCCT*CCATGTACCGGCAAAAGCAAGTGTAGC | Handle staple for Cy3_66 |
| *CTCCTATCTCCAATCACTCCT*CAAACTACAGGCGAAAACCACACCCGC | Handle staple for Cy3_67 |
| *CTCCTATCTCCAATCACTCCT*CCTCATAGTTGCAGCACAGGGCGCGTA | Handle staple for Cy3_68 |
| *CTCCTATCTCCAATCACTCCT*GTCTTTCCGCTGATTGCACGTATAACG | Handle staple for Cy3_69 |
| *CTCCTATCTCCAATCACTCCT*TGGGATTTGTGGTTTTAGAGCGGGAGC | Handle staple for Cy3_70 |
| *CTCCTATCTCCAATCACTCCT*CGGAGTGAGCGGGGAGAGGGATTTTAG | Handle staple for Cy3_71 |
| *CTCCTATCTCCAATCACTCCT*TGCGAATAGTCGTGCCATCCTGAGAAG | Handle staple for Cy3_72 |
| *CTCCTATCTCCAATCACTCCT*AAAAAAAATGCGCTCAGCCACCGAGTA | Handle staple for Cy3_73 |
| *CTCCTATCTCCAATCACTCCT*TCGGTTTAGCCTAATGCAAATTAACCG | Handle staple for Cy3_74 |

| | |
|---|---|
| *CTCCTATCTCCAATCACTCCT*TAAACAGCACGAGCCGATTAGTAATAA | Handle staple for Cy3_75 |
| *CTCCTATCTCCAATCACTCCT*GACAACAAGAAATTGTAGAACTCAAAC | Handle staple for Cy3_76 |
| *CTCCTATCTCCAATCACTCCT*TCGGTCGCAATTCGTAATCCAGAACAA | Handle staple for Cy3_77 |
| *CTCCTATCTCCAATCACTCCT*CTTTTGCGCCTCACAGAACAGGAAAAA | Handle staple for Cy3_78 |
| *CTCCTATCTCCAATCACTCCT*CAGCATCGAGCACGCGATTTTGACGCT | Handle staple for Cy3_79 |
| *CTCCTATCTCCAATCACTCCT*GCTTTGAGGGCCGTTTATTTACATTGG | Handle staple for Cy3_80 |
| *CTCCTATCTCCAATCACTCCT*TTCCATTAGTGCACTCACCAGTAATAA | Handle staple for Cy3_81 |
| *CTCCTATCTCCAATCACTCCT*CGAAGGCAATCAGACGAGAGATAGAAC | Handle staple for Cy3_82 |
| *CTCCTATCTCCAATCACTCCT*AATACACTGTTACCTGAAGAATACGTG | Handle staple for Cy3_83 |
| *CTCCTATCTCCAATCACTCCT*ATTATACCTGTTCAGCATGGCTATTAG | Handle staple for Cy3_84 |
| *CTCCTATCTCCAATCACTCCT*TTTGTATCTCTTTGCTTAGCCCTAAAA | Handle staple for Cy3_85 |
| *CTCCTATCTCCAATCACTCCT*CGCGACCTACTGTTGCGAACGAACCAC | Handle staple for Cy3_86 |
| *CTCCTATCTCCAATCACTCCT*GCGCAGACATCCGCCGGGTGAGGCGGT | Handle staple for Cy3_87 |
| *CTCCTATCTCCAATCACTCCT*CCAACTTTATCAGCGGAACAGTGCCAC | Handle staple for Cy3_88 |
| *CTCCTATCTCCAATCACTCCT*GACCAGGCCCACGCAATGAAAAATCTA | Handle staple for Cy3_89 |
| *CTCCTATCTCCAATCACTCCT*AGTAATCTGAATGCCACTCAAATATCA | Handle staple for Cy3_90 |
| *CTCCTATCTCCAATCACTCCT*AAATCAACAGCGTGGTGTCAGTTGGCA | Handle staple for Cy3_91 |
| *CTCCTATCTCCAATCACTCCT*AAGGCTTGCTCATAACTTGAGGAAGGT | Handle staple for Cy3_92 |
| *CTCCTATCTCCAATCACTCCT*AGTAAATTTCTCGTCGAGCACTAACAA | Handle staple for Cy3_93 |
| *CTCCTATCTCCAATCACTCCT*TTAATCATATTGCCGTAATAGATAATA | Handle staple for Cy3_94 |
| *CTCCTATCTCCAATCACTCCT*AACTGGCTCTTTAGTGATTAGACTTTA | Handle staple for Cy3_95 |
| *CTCCTATCTCCAATCACTCCT*GAAAAATCAAAAAAATTATTAAATCCT | Handle staple for Cy3_96 |
| *CTCCTATCTCCAATCACTCCT*CAACATTACCGCCAGCTTTAAAAGTTT | Handle staple for Cy3_97 |
| *CTCCTATCTCCAATCACTCCT*AGATTTAGAACAGCGGCGGAACAAAGA | Handle staple for Cy3_98 |
| *CTCCTATCTCCAATCACTCCT*ACATAACGGAAGGGATATTATCATCAT | Handle staple for Cy3_99 |
| *CTCCTATCTCCAATCACTCCT*GAGCAACACCGGAATTTGGCAATTCAT | Handle staple for Cy3_100 |
| *CTCCTATCTCCAATCACTCCT*ACGATAAAATCGGCGATGGATTATACT | Handle staple for Cy3_101 |
| *CTCCTATCTCCAATCACTCCT*AAAGAAGTGCCGCCACAGAACCTACCA | Handle staple for Cy3_102 |
| *CTCCTATCTCCAATCACTCCT*GTTTAGACAAACGACGTAAAACAGAAA | Handle staple for Cy3_103 |
| *CTCCTATCTCCAATCACTCCT*GTCATAAAGTAACGCCTTCAGGTTTAA | Handle staple for Cy3_104 |
| *CTCCTATCTCCAATCACTCCT*TTAAACAGGAAAGGGGAACAGTACCTT | Handle staple for Cy3_105 |
| *CTCCTATCTCCAATCACTCCT*CAAAAATCCGGTGCGGAACGGATTCGC | Handle staple for Cy3_106 |
| *CTCCTATCTCCAATCACTCCT*AGAAGCAACCATTCAGAGTTACAAAAT | Handle staple for Cy3_107 |

| | |
|---|---|
| *CTCCTATCTCCAATCACTCCT*GGAAGCCCTTCTGGTGCATTTCAATTA | Handle staple for Cy3_108 |
| *CTCCTATCTCCAATCACTCCT*TTCGAGCTGAAGATCGTGAAACAAACA | Handle staple for Cy3_109 |
| *CTCCTATCTCCAATCACTCCT*CTCCAACACAGTTTGATACATTTAACA | Handle staple for Cy3_110 |
| *CTCCTATCTCCAATCACTCCT*TGCTCCTTGGTGTAGATTTTAATGGAA | Handle staple for Cy3_111 |
| *CTCCTATCTCCAATCACTCCT*GCTTAGAGACGGCGGAATGTGAGTGAA | Handle staple for Cy3_112 |
| *CTCCTATCTCCAATCACTCCT*CTCAACATGAGTAACACGTCGCTATTA | Handle staple for Cy3_113 |
| *CTCCTATCTCCAATCACTCCT*TGTCTGGACTGTAGCCCCTTGAAAACA | Handle staple for Cy3_114 |
| *CTCCTATCTCCAATCACTCCT*CCCAATTCGAACGCCAGACGCTGAGAA | Handle staple for Cy3_115 |
| *CTCCTATCTCCAATCACTCCT*ATTAGATATTTTTGTTCAAAATCATAG | Handle staple for Cy3_116 |
| *CTCCTATCTCCAATCACTCCT*TTAGCTATTAAACGTTTAACCTCCGGC | Handle staple for Cy3_117 |
| *CTCCTATCTCCAATCACTCCT*AGGTGGCAAACAGGAATATATGTAAAT | Handle staple for Cy3_118 |
| *CTCCTATCTCCAATCACTCCT*AACATCCAATGTACCCCAAGACAAAGA | Handle staple for Cy3_119 |
| *CTCCTATCTCCAATCACTCCT*TTAGCAAATGAACGGTAATATATTTTA | Handle staple for Cy3_120 |
| *CTCCTATCTCCAATCACTCCT*AAGCTAAATTGCCTGACTAAATTTAAT | Handle staple for Cy3_121 |
| *CTCCTATCTCCAATCACTCCT*CTGTAATATATTTTTGGTGATAAATAA | Handle staple for Cy3_122 |
| *CTCCTATCTCCAATCACTCCT*ACGCAAGGTTCTAGCTCACCGGAATCA | Handle staple for Cy3_123 |
| *CTCCTATCTCCAATCACTCCT*TTTTAAATACAGTCAAGTTTAGTATCA | Handle staple for Cy3_124 |

**Table S4**

Antihandle carrying Cy3 to label the NS.

| **Sequence (5’ to 3’)** | **Name** |
|---|---|
| Cy3/AGGAGTGATTGGAGATAGGAG | Cy3-labeled antihandle |

**Table S5**

Core staples to build the DNA calibration rod.

| **Sequence (5' to 3')** | **Name** |
|---|---|
| AACGATCTAAAGTAAAGCCGGATTAAAGAACGTGGAC | Core Staple_01 |
| AGCATTCCACAGAAGAAAGCGAGTGTTGTTCCAGTTT | Core Staple_02 |
| GTTTCGTCACCAGGCAAGTGTATAAATCAAAAGAATA | Core Staple_03 |
| GCCCAATAGGAACCACACCCGTGATGGTGGTTCCGAA | Core Staple_04 |
| AGAGCCACCACCCGCGCGTACGCTGGTTTGCCCCAGC | Core Staple_05 |
| TCAGAACCGCCACTAACGTGCTGGCCCTGAGAGAGTT | Core Staple_06 |
| CACCGTACTCAGGAGCTAAACAGACGGGCAACAGCTG | Core Staple_07 |
| GTTGATATAAGTAGACAGGAAGGGCGCCAGGGTGGTT | Core Staple_08 |
| TCAGTACCAGGCGGTTTTTATGCCAACGCGCGGGGAG | Core Staple_09 |
| CAAGAGAAGGATTGAGTCTGTAAACCTGTCGTGCCAG | Core Staple_10 |
| AAACATGAAAGTAGCAATACTGCGTTGCGCTCACTGC | Core Staple_11 |
| GCCCCCTGCCTATTGCCTGAGGTGCCTAATGAGTGAG | Core Staple_12 |
| CTTGAGTAACAGTTGCTGGTAACGAGCCGGAAGCATA | Core Staple_13 |
| GTACTGGTAATAAGCCATTGCAATTGTTATCCGCTCA | Core Staple_14 |
| GCGTCATACATGGACCTACATCGTAATCATGGTCATA | Core Staple_15 |
| AAGCGCAGTCTCTGATTATTTAGTTGAGGATCCCCGG | Core Staple_16 |
| AACAAATAAATCCCGACCAGTTGCCTGTTCTTCGCGT | Core Staple_17 |
| GGCAGGTCAGACGAGAGATAGACGGTCATACCGGGGG | Core Staple_18 |
| CAGAGCCGCCGCCGAATACGTGTGCTGCGGCCAGAAT | Core Staple_19 |
| ACCACCCTCAGAGCTATTAGTCGCAGTGTCACTGCGC | Core Staple_20 |
| CCGCCACCCTCAGTAAAACATGGTGCCGGTGCCCCCT | Core Staple_21 |
| CAGAGCCACCACCCACCAGCACGGCATCAGATGCCGG | Core Staple_22 |
| TGCCATCTTTTCATCAGTATTCCCTTACACTGGTGTG | Core Staple_23 |
| GCATTTTCGGTCACTGAGAGCTGGGTAAAGGTTTCTT | Core Staple_24 |
| CTTTAGCGTCAGACATCACCTGAGCCGGGTCACTGTT | Core Staple_25 |
| ACCGTAATCAGTATCAATCAACGCACTCAATCCGCCG | Core Staple_26 |
| GAAACGTCACCAAAGTTGAAATCCAGCATCAGCGGGG | Core Staple_27 |
| CACCAGTAGCACCTATCTTTACATCCCACGCAACCAG | Core Staple_28 |
| TTGAGCCATTTGGGAGCCGTCAAGAATGCCAACGGCA | Core Staple_29 |
| TTCATTAAAGGTGAGAAGTATAGCGTGGTGCTGGTCT | Core Staple_30 |
| GAGGGAGGGAAGGCTCGTATTCATAACGGAACGTGCC | Core Staple_31 |

| | |
|---|---|
| GCCAAAGACAAAAATTTTAAAGTCGCTGGCAGCCTCC | Core Staple_32 |
| CAATCAATAGAAACGGAACAAGTTCCGGCAAACGCGG | Core Staple_33 |
| CAAAGACACCACGTATCATCAATGAAGGGTAAAGTTA | Core Staple_34 |
| ATACATAAAGGTGAATTCATCCGTAAAAAAAGCCGCA | Core Staple_35 |
| TTACGCAGTATGTATACTTCTGGGCGGTTGTGTACAT | Core Staple_36 |
| CCCAAAAGAACTGCCATATCACTTAAATTTCTGCTCA | Core Staple_37 |
| GGAAACCGAGGAAATAAAGAACGGAAAAAGAGACGCA | Core Staple_38 |
| AGTAAGCAGATAGGTCAGATGAGACTTTCTCCGTGGT | Core Staple_39 |
| ATAGCTATCTTACCATCGGGACATGTTTACCAGTCCC | Core Staple_40 |
| CAATAATAAGAGCTGCTTTGATCACCGGAAACAATCG | Core Staple_41 |
| AGAGAGATAACCCAGGCGAATCAGAGGTGGAGCCGCC | Core Staple_42 |
| ACAAAGTCAGAGGAAGAAGATACGTTGTAAAACGACG | Core Staple_43 |
| CATTAGACGGGAGAAATTAATAGTTGGGTAACGCCAG | Core Staple_44 |
| TACAGAGAGAATATACCTTTTTGGCGAAAGGGGGATG | Core Staple_45 |
| TGTTTAACGTCAAATATATGTATCGGTGCGGGCCTCT | Core Staple_46 |
| ATTTATCCCAATCATCGTCGCCCATTCAGGCTGCGCA | Core Staple_47 |
| AATTTGCCAGTTACCTTGAAACTGGTGCCGGAAACCA | Core Staple_48 |
| TACCAACGCTAACCGCTGAGAATCGCACTCCAGCCAG | Core Staple_49 |
| TTTGCACCCAGCTATCATAGGGAGGGGACGACGACAG | Core Staple_50 |
| TTTTGAAGCCTTACCGGCTTATGGGCGCATCGTAACC | Core Staple_51 |
| GCGTTTTAGCGAAAATGCTGAGACCGTAATGGGATAG | Core Staple_52 |
| AGAAGGCTTATCCAACGCGAGGTCGGATTCTCCGTGG | Core Staple_53 |
| TTACCGCGCCCAATTAATTTCCATCAACATTAAATGT | Core Staple_54 |
| GCAAGCCGTTTTTTTGAAATAAATTCGCGTCTGGCCT | Core Staple_55 |
| ATTAAACCAAGTATAAATAAGATTTTTTAACCAATAG | Core Staple_56 |
| GGCTGTCTTTCCTTAGAAAAAAAATTCGCATTAAATT | Core Staple_57 |
| ATTTACGAGCATGTACAAATTATATTTAAATTGTAAA | Core Staple_58 |
| GTCCTGAACAAGAAACAGTAGAAGCCCCAAAAACAGG | Core Staple_59 |
| GCAGAACGCGCCTTAACAACGTCAATCATATGTACCC | Core Staple_60 |
| GACGACGACAATAATTTTCGAAATCGATGAACGGTAA | Core Staple_61 |
| GAGCTTGACGGGGTTTGTCGTCAACAGTTTCAGCGGA | Core Staple_62 |
| AGAAAGGAAGGGACAGCCCTCACAACTAAAGGAATTG | Core Staple_63 |
| CGCTAGGGCGCTGTACAAACTCACGTTGAAAATCTCC | Core Staple_64 |

| | |
|---|---|
| TGCGCGTAACCACCCATGTACAAGGAGCCTTTAATTG | Core Staple_65 |
| GCGCCGCTACAGGTCATTTTCTGCTTTCGAGGTGAAT | Core Staple_66 |
| TGACGAGCACGTACCTCAGAATACCGATAGTTGCGCC | Core Staple_67 |
| GAATCAGAGCGGGAGGTTTAGATCGCCCACGCATAAC | Core Staple_68 |
| TAAAGGGATTTTATAGCCCGGTGAGGCTTGCAGGGAG | Core Staple_69 |
| AATCCTGAGAAGTGATAAGTGCGGGATCGTCACCCTC | Core Staple_70 |
| CCACCGAGTAAAAAGGATTAGATCGGAACGAGGGTAG | Core Staple_71 |
| ATTAACCGTTGTATTAAGAGGTTTGAGGACTAAAGAC | Core Staple_72 |
| TAATAACATCACTTTCGGAACTTCCATTAAACGGGTA | Core Staple_73 |
| AAACTATCGGCCTGCCCGTATTACGAAGGCACCAACC | Core Staple_74 |
| AATATTACCGCCAGTTTTAACAAAGAATACACTAAAA | Core Staple_75 |
| CGCTCATGGAAATCTTTTGATCCAGCGATTATACCAA | Core Staple_76 |
| ATCGTCTGAAATGGAATTTACAACGGAGATTTGTATC | Core Staple_77 |
| TTCACCAGTCACATCATTAAATGTCGAAATCCGCGAC | Core Staple_78 |
| CATTCTGGCCAACATTGGCCTAGCCGGAACGAGGCGC | Core Staple_79 |
| CCTGAAAGCGTAAAGCATTGAGGGAACCGAACTGACC | Core Staple_80 |
| TATTTTTGAATGGCCGCCACCAGATGAACGGTGTACA | Core Staple_81 |
| GAACTGATAGCCCAACCGCCATGGCTGACCTTCATCA | Core Staple_82 |
| ATACCGAACGAACGGAACCGCGAACCGGATATTCATT | Core Staple_83 |
| AGAGGTGAGGCGGTAATCAAACAAAGCTGCTCATTCA | Core Staple_84 |
| CAACAGTGCCACGTAGCCCCCCTGACGAGAAACACCA | Core Staple_85 |
| GAAAAATCTAAAGCTGTAGCGGGGCTTGAGATGGTTT | Core Staple_86 |
| AAATATCAAACCCGCGACAGAATTGTGAATTACCTTA | Core Staple_87 |
| TTGGCAAATCAACTGAAACCAGGCTCATTATACCAGT | Core Staple_88 |
| AGGTTATCTAAAAATTACCATAAAATCTACGTTAATA | Core Staple_89 |
| AACTAATAGATTAGAATTAGACAACATTATTACAGGT | Core Staple_90 |
| CATTTGAGGATTTAATTATCATGAGATTTAGGAATAC | Core Staple_91 |
| AACAATTCGACAATAAATATTAGATACATAACGCCAA | Core Staple_92 |
| CCGAACGTTATTAGGGCGACATAGTAAGAGCAACACT | Core Staple_93 |
| CATTATCATTTTGATTCATATACCAGACGACGATAAA | Core Staple_94 |
| GAAGGAGCGGAATGAATAAGTGGCTTTTGCAAAAGAA | Core Staple_95 |
| ATCAGATGATGGCGCAACATATAATAGTAAAATGTTT | Core Staple_96 |
| GATTGTTTGGATTTAGCAAACAATACTGCGGAATCGT | Core Staple_97 |

| | |
|---|---|
| GGGTTAGAACCTAGCATGATTATCCCCCTCAAATGCT | Core Staple_98 |
| ACGTAAAACAGAAACGCAATAACGAGAATGACCATAA | Core Staple_99 |
| TTTCAGGTTTAACCCGAACAATTACCCTGACTATTAT | Core Staple_100 |
| ACAGTACCTTTTACGAAGCCCGATTGCATCAAAAAGA | Core Staple_101 |
| GGATTCGCCTGATAAGAAACAAAGACTTCAAATATCG | Core Staple_102 |
| CAAAATCGCGCAGACAAGAATTCAAAGCGAACCAGAC | Core Staple_103 |
| ATTACCTGAGCAAGTAATTGACAGGTCAGGATTAGAG | Core Staple_104 |
| CATCAAGAAAACAAATTAACTCCTTTTGATAAGAGGT | Core Staple_105 |
| ATTTCATTTGAATACATAAAATTAGAGCTTAATTGCT | Core Staple_106 |
| AGTACATAAATCAAAATGAAACTCAACATGTTTTAAA | Core Staple_107 |
| CTTGCTTCTGTAACAAATAAGGGTGTCTGGAAGTTTC | Core Staple_108 |
| TTTCCCTTAGAATCAAAATAAGATTCCCAATTCTGCG | Core Staple_109 |
| GCTTAGATTAAGAGAGCGTCTTTGACCATTAGATACA | Core Staple_110 |
| TGAATTTATCAAAACAATTTTTAACCTGTTTAGCTAT | Core Staple_111 |
| ACCTTTTTAACCTAATCAAGACGAGCTGAAAAGGTGG | Core Staple_112 |
| ATAACTATATGTACCTCCCGAAGTAGTAGCATTAACA | Core Staple_113 |
| TCGCAAGACAAAGGGTATTCTGGCAAGGCAAAGAATT | Core Staple_114 |
| AAATATATTTTAGTAGCAAGCAAAGCCTCAGAGCATA | Core Staple_115 |
| TAAATTTAATGGTATTTTCATACCAAAAACATTATGA | Core Staple_116 |
| ATAAATAAGGCGTCCGCACTCCGGGAGAAGCCTTTAT | Core Staple_117 |
| GAATCATAATTACTATCATTCAAATTTTTAGAACCCT | Core Staple_118 |
| ATCATATGCGTTATAGAAACCAATGCCTGAGTAATGT | Core Staple_119 |
| AAAGCCAACGCTCAAAATAATAAGGGTGAGAAAGGCC | Core Staple_120 |
| GAATCGCCATATTGTTTATCACCATCAATATGATATT | Core Staple_121 |
| TTTAGGCAGAGGCAACAACATTAAATTAATGCCGGAG | Core Staple_122 |
| GAAACAGCTTCAGAAAATAACGGAATA | 398_501nm_Cy3nega_1 |
| GAAGGGATTCAGGTCTAGTTACCAGAA | 398_501nm_Cy3nega_2 |
| GGAATTTGGCAAAGCGTTTTTAAGAAA | 398_501nm_Cy3nega_3 |
| ACGGGAACTTCGAGCTTGAGTTAAGCC | 398_501nm_Cy3nega_4 |
| GCCAGTGCAACTCCAAGCGCTAATATC | 398_501nm_Cy3nega_5 |
| GGTTTTCCTAATTGCTGAACACCCTGA | 398_501nm_Cy3nega_6 |
| TGCTGCAACGGATGGCACAGGGAAGCG | 398_501nm_Cy3nega_7 |
| TCGCTATTTGCTGTAGATAGCAGCCTT | 398_501nm_Cy3nega_8 |

| | |
|---|---|
| ACTGTTGGTAAAGTACAAACGATTTTT | 398_501nm_Cy3nega_9 |
| ATCAAAAAAGCTCTCAATTGCGTAGAT | 398_501nm_Cy3nega_10 |
| TTAAGAGGTACAGCGCGAAACAATAAC | 398_501nm_Cy3nega_11 |
| CGTTTTAAGGATAACCATACCAAGTTA | 398_501nm_Cy3nega_12 |
| CGGAAGCACAAGCTTTTATTCATTTCA | 398_501nm_Cy3nega_13 |
| AGTACCTTCAGTCACGGATGAAACAAA | 398_501nm_Cy3nega_14 |
| CATTTTTGGGCGATTATACATTTAACA | 398_501nm_Cy3nega_15 |
| GAATATAAACGCCAGCTTAATGGAAAC | 398_501nm_Cy3nega_16 |
| TATGCAACGAAGGGCGGAGTGAATAAC | 398_501nm_Cy3nega_17 |
| GGCAAAGCTAACAGTTACAGCCATATT | 501_599nm_Cy3nega_1 |
| CTTTCCGGGATTTAGTTTCCAGAGCCT | 501_599nm_Cy3nega_2 |
| TATCGGCCATGGTCAAATCCTGAATCT | 501_599nm_Cy3nega_3 |
| GTGCATCTTTGGGGCGTTAGTTGCTAT | 501_599nm_Cy3nega_4 |
| GTCACGTTCTACTAATCTTGCGGGAGG | 501_599nm_Cy3nega_5 |
| GAACAAACATCATACAAAGAACGCGAG | 501_599nm_Cy3nega_6 |
| GAGCGAGTTAAGCAATAAATCAGATAT | 501_599nm_Cy3nega_7 |
| TCCTGTAGTCGGTTGTCGTAGGAATCA | 501_599nm_Cy3nega_8 |
| GAACGCCATACTTTTGATCGAGAACAA | 501_599nm_Cy3nega_9 |
| ATTCCATAGCCATTCGTATTAATTAAT | 501_599nm_Cy3nega_10 |
| AACGAGTACACCGCTTACATAGCGATA | 501_599nm_Cy3nega_11 |
| TTTCGCAATCAGGAAGAGAGTCAATAG | 501_599nm_Cy3nega_12 |
| ATTTTCATGCCAGTTTTCTGAGAGACT | 501_599nm_Cy3nega_13 |
| CATCAATTGGTGTAGAGGTTGGGTTAT | 501_599nm_Cy3nega_14 |
| TCCAATAAGGCGGATTTGCAAATCCAA | 501_599nm_Cy3nega_15 |
| AGCAAAATAACAACCCAAAACTTTTTC | 501_599nm_Cy3nega_16 |
| AAGCTAAACCAGCTTTATCTTCTGACC | 501_599nm_Cy3nega_17 |
| CCCTGTAATCAAAAATCCGACCGTGTG | 501_599nm_Cy3nega_18 |
| TTTGTTAAAAGGATAACAAGAACGGGT | 599_658nm_Cy3nega_1 |
| CGTTAATATTAAATGCAATCAATAATC | 599_658nm_Cy3nega_2 |
| AAGATTGTAGATTCAAATCCCATCCTA | 599_658nm_Cy3nega_3 |
| CGGTTGATTCAAATCAACAATAGATAA | 599_658nm_Cy3nega_4 |
| TCGTAAAACTAGCTGAGTTCAGCTAAT | 599_658nm_Cy3nega_5 |
| TTCAACGCATCAGCTCAATAAACACCG | 599_658nm_Cy3nega_6 |

| | |
|---|---|
| CATATATTTTTTGTTAGCCTGTTTAGT | 599_658nm_Cy3nega_7 |
| GTAGGTAAATAAGCAACTTACCAGTAT | 599_658nm_Cy3nega_8 |
| GGAGACAGAATCAGAAGGCTTAATTGA | 599_658nm_Cy3nega_9 |
| AGGGTAGCAACAAGAGGCCAGTAATAA | 599_658nm_Cy3nega_10 |

**Table S6**

Handle staples for the DNA calibration rod. Sequences in italics indicate the handle site.

Handle staples for biotin

| **Sequence (5' to 3')** | **Name** |
|---|---|
| GGAACAAGAGAAAGGAATAGTTAGCGT *CTCTCCTCTCCACCATATCCA* | Handle for biotin #1 |
| GTTACCTGCTTGACAACTCCCTCAGAG *CTCTCCTCTCCACCATATCCA* | Handle for biotin #2 |
| GCGAAACGAAGCCCGAATGAAATAGCA *CTCTCCTCTCCACCATATCCA* | Handle for biotin #3 |
| CTGGAGCATATTTTTGAATTCTGTCCA *CTCTCCTCTCCACCATATCCA* | Handle for biotin #4 |
| TTTGCTAACAAAGGGCCCCCCGATTTA *CTCTCCTCTCCACCATATCCA* | Handle for biotin #5 |
| GACCAGGCCGATCCAGCTTTAATGCGC *CTCTCCTCTCCACCATATCCA* | Handle for biotin #6 |
| AGTCAGAATGAGAGATAATATACAGTA *CTCTCCTCTCCACCATATCCA* | Handle for biotin #7 |
| CAACCGTTCTAGCATGCCAACATGTAA *CTCTCCTCTCCACCATATCCA* | Handle for biotin #8 |

Handle staples for Cy3 fluorophore in a 398-nm long DNA calibration rod.

| **Sequence (5' to 3')** | **Name** |
|---|---|
| *CTCCTATCTCCAATCACTCCT*TCCAACGTACAACTTTCTTTCCAGACG | Handle staple for Cy3_398nm rod_1 |
| *CTCCTATCTCCAATCACTCCT*GCCCGAGAAATTTTTTACAACGCCTGT | Handle staple for Cy3_398nm rod_2 |
| *CTCCTATCTCCAATCACTCCT*ATCGGCAAGGCTCCAACGTAACACTGA | Handle staple for Cy3_398nm rod_3 |
| *CTCCTATCTCCAATCACTCCT*AGGCGAAATATCAGCTAGGGATAGCAA | Handle staple for Cy3_398nm rod_4 |

| | |
|---|---|
| *CTCCTATCTCCAATCACTCCT*GCAGCAAGCAGCTTGACCGCCACCCTC | Handle staple for Cy3_398nm rod_5 |
| *CTCCTATCTCCAATCACTCCT*ATTGCCCTCAACAACCTACCGCCACCC | Handle staple for Cy3_398nm rod_6 |
| *CTCCTATCTCCAATCACTCCT*TTTCTTTTTCGGTCGCAATAGGTGTAT | Handle staple for Cy3_398nm rod_7 |
| *CTCCTATCTCCAATCACTCCT*AGGCGGTTCGCTTTTGCCGTCGAGAGG | Handle staple for Cy3_398nm rod_8 |
| *CTCCTATCTCCAATCACTCCT*CTGCATTAAAGACAGCCGGGGTTTTGC | Handle staple for Cy3_398nm rod_9 |
| *CTCCTATCTCCAATCACTCCT*CCGCTTTCACAGAGGCCTGAGACTCCT | Handle staple for Cy3_398nm rod_10 |
| *CTCCTATCTCCAATCACTCCT*CTAACTCAGAGGAAGTCTATTATTCTG | Handle staple for Cy3_398nm rod_11 |
| *CTCCTATCTCCAATCACTCCT*AAGTGTAAAATGCCACAAACAGTTAAT | Handle staple for Cy3_398nm rod_12 |
| *CTCCTATCTCCAATCACTCCT*CAATTCCAAAGAGGCAGGGGTCAGTGC | Handle staple for Cy3_398nm rod_13 |
| *CTCCTATCTCCAATCACTCCT*GCTGTTTCTTTGACCCGATACAGGAGT | Handle staple for Cy3_398nm rod_14 |
| *CTCCTATCTCCAATCACTCCT*GTACCGAGCAAAGTACCGTTCCAGTAA | Handle staple for Cy3_398nm rod_15 |
| *CTCCTATCTCCAATCACTCCT*CCGTGAGCATAAATTGGCCAGAATGGA | Handle staple for Cy3_398nm rod_16 |
| *CTCCTATCTCCAATCACTCCT*TTTCTGCCTGTTACTTTGATATTCACA | Handle staple for Cy3_398nm rod_17 |
| *CTCCTATCTCCAATCACTCCT*GCGGCGGGAATCATAACAGGAGGTTGA | Handle staple for Cy3_398nm rod_18 |
| *CTCCTATCTCCAATCACTCCT*GCCTGTGCAAGAGGACAGAACCACCAC | Handle staple for Cy3_398nm rod_19 |
| *CTCCTATCTCCAATCACTCCT*GCATCAGAGCATAGGCCCCTCAGAGCC | Handle staple for Cy3_398nm rod_20 |
| *CTCCTATCTCCAATCACTCCT*TTCAGCAACAACGTAAATCACCGGAAC | Handle staple for Cy3_398nm rod_21 |
| *CTCCTATCTCCAATCACTCCT*TGCTCGTCGGCTTGCCTTATTAGCGTT | Handle staple for Cy3_398nm rod_22 |
| *CTCCTATCTCCAATCACTCCT*GCCCTGCGAGTAAATTCGTTTTCATCG | Handle staple for Cy3_398nm rod_23 |
| *CTCCTATCTCCAATCACTCCT*GGCGCGGTCTTTAATCATCAAGTTTGC | Handle staple for Cy3_398nm rod_24 |
| *CTCCTATCTCCAATCACTCCT*TCATTGCATAAGAACTTCGATAGCAGC | Handle staple for Cy3_398nm rod_25 |
| *CTCCTATCTCCAATCACTCCT*CTTACGGCTGGGAAGATAGCAAGGCCG | Handle staple for Cy3_398nm rod_26 |

| | |
|---|---|
| *CTCCTATCTCCAATCACTCCT*GCACCGTCTAACGGAAGCCAGCAAAAT | Handle staple for Cy3_398nm rod_27 |
| *CTCCTATCTCCAATCACTCCT*GGTCAGCATCATCAGTCCGTCACCGAC | Handle staple for Cy3_398nm rod_28 |
| *CTCCTATCTCCAATCACTCCT*GGACTTGTACTAATGCGACGGAAATTA | Handle staple for Cy3_398nm rod_29 |
| *CTCCTATCTCCAATCACTCCT*GGCCAGAGACGAGGCATTCAACCGATT | Handle staple for Cy3_398nm rod_30 |
| *CTCCTATCTCCAATCACTCCT*TCCGTTTTCCTCGTTTGGTTTACCAGC | Handle staple for Cy3_398nm rod_31 |
| *CTCCTATCTCCAATCACTCCT*AACGATGCTAGCGAGATTATTTTGTCA | Handle staple for Cy3_398nm rod_32 |
| *CTCCTATCTCCAATCACTCCT*CAGGCGGCAGAGGGGGTAAAAGAAACG | Handle staple for Cy3_398nm rod_33 |
| *CTCCTATCTCCAATCACTCCT*CGACATAATAGCGTCCGTAGAAAATAC | Handle staple for Cy3_398nm rod_34 |
| *CTCCTATCTCCAATCACTCCT*TTTGCCGCTTCATTGAAAGACTCCTTA | Handle staple for Cy3_398nm rod_35 |
| *CTCCTATCTCCAATCACTCCT*GTGAGAATAGTCCACTCGAACGTGGCG | Handle staple for Cy3_398nm rod_36 |
| *CTCCTATCTCCAATCACTCCT*CGAATAATTAGGGTTGAAAGGAGCGGG | Handle staple for Cy3_398nm rod_37 |
| *CTCCTATCTCCAATCACTCCT*AAAAAAAAAATCCCTTAGCGGTCACGC | Handle staple for Cy3_398nm rod_38 |
| *CTCCTATCTCCAATCACTCCT*TATCGGTTATCCTGTTCCGCGCTTAAT | Handle staple for Cy3_398nm rod_39 |
| *CTCCTATCTCCAATCACTCCT*TTCTTAAACGGTCCACTATGGTTGCTT | Handle staple for Cy3_398nm rod_40 |
| *CTCCTATCTCCAATCACTCCT*GACAATGATCACCGCCTTTCCTCGTTA | Handle staple for Cy3_398nm rod_41 |
| *CTCCTATCTCCAATCACTCCT*CGATATATCACCAGTGAGGAGGCCGAT | Handle staple for Cy3_398nm rod_42 |
| *CTCCTATCTCCAATCACTCCT*TTAAAGGCTGCGTATTCGGTACGCCAG | Handle staple for Cy3_398nm rod_43 |
| *CTCCTATCTCCAATCACTCCT*AGCAGCGAATGAATCGAATCAGTGAGG | Handle staple for Cy3_398nm rod_44 |
| *CTCCTATCTCCAATCACTCCT*CAACGGCTCAGTCGGGCCATCACGCAA | Handle staple for Cy3_398nm rod_45 |
| *CTCCTATCTCCAATCACTCCT*TTTTTCATCATTAATTTCTTTGATTAG | Handle staple for Cy3_398nm rod_46 |
| *CTCCTATCTCCAATCACTCCT*AAATACGTAGCCTGGGTAGAAGAACTC | Handle staple for Cy3_398nm rod_47 |
| *CTCCTATCTCCAATCACTCCT*TAAAACGACACAACATATATCCAGAAC | Handle staple for Cy3_398nm rod_48 |

| | |
|---|---|
| *CTCCTATCTCCAATCACTCCT*CACTCATCCTGTGTGAAACAGGAAAAA | Handle staple for Cy3_398nm rod_49 |
| *CTCCTATCTCCAATCACTCCT*GCGCGAAACTCGAATTTTTGACGCTCA | Handle staple for Cy3_398nm rod_50 |
| *CTCCTATCTCCAATCACTCCT*ATCGCCTGCTCCTCACACATTGGCAGA | Handle staple for Cy3_398nm rod_51 |
| *CTCCTATCTCCAATCACTCCT*CTGCTCCAAGCACGCGAATAAAAGGGA | Handle staple for Cy3_398nm rod_52 |
| *CTCCTATCTCCAATCACTCCT*AGACGGTCCCGTTTTCAACCCTTCTGA | Handle staple for Cy3_398nm rod_53 |
| *CTCCTATCTCCAATCACTCCT*AACTTTGAACTCTGTGGGCACAGACAA | Handle staple for Cy3_398nm rod_54 |
| *CTCCTATCTCCAATCACTCCT*AGAGTAATCAGCCAGCCGCCATTAAAA | Handle staple for Cy3_398nm rod_55 |
| *CTCCTATCTCCAATCACTCCT*ACCCAAATATCGTTAAGAAGATAAAAC | Handle staple for Cy3_398nm rod_56 |
| *CTCCTATCTCCAATCACTCCT*GTGAATAAATAAACATAACACCGCCTG | Handle staple for Cy3_398nm rod_57 |
| *CTCCTATCTCCAATCACTCCT*GAACGAGTGCTGGTAACAGCAGCAAAT | Handle staple for Cy3_398nm rod_58 |
| *CTCCTATCTCCAATCACTCCT*AATTTCAATGCGGTATTGCTGAACCTC | Handle staple for Cy3_398nm rod_59 |
| *CTCCTATCTCCAATCACTCCT*TGCGATTTGGCGCTTTTATCTGGTCAG | Handle staple for Cy3_398nm rod_60 |
| *CTCCTATCTCCAATCACTCCT*CAGGACGTTGGAGGTGGGAATTGAGGA | Handle staple for Cy3_398nm rod_61 |
| *CTCCTATCTCCAATCACTCCT*AAACGAACGGTGGTGCGGAGCACTAAC | Handle staple for Cy3_398nm rod_62 |
| *CTCCTATCTCCAATCACTCCT*AGAAAGATGCAACCGCAATAGATAATA | Handle staple for Cy3_398nm rod_63 |
| *CTCCTATCTCCAATCACTCCT*CACATTCAAGAACGTCTAGACTTTACA | Handle staple for Cy3_398nm rod_64 |
| *CTCCTATCTCCAATCACTCCT*AAGGAATTCACATCCTAAATCCTTTGC | Handle staple for Cy3_398nm rod_65 |
| *CTCCTATCTCCAATCACTCCT*ATCATAACTTCGTCTCAGTTTGAGTAA | Handle staple for Cy3_398nm rod_66 |
| *CTCCTATCTCCAATCACTCCT*AACCAAAATGATTGCCAGAAACCACCA | Handle staple for Cy3_398nm rod_67 |
| *CTCCTATCTCCAATCACTCCT*GTTTTGCCCTTTAGTGTATTCCTGATT | Handle staple for Cy3_398nm rod_68 |
| *CTCCTATCTCCAATCACTCCT*AGACTGGAAAAAATCCAATATAATCCT | Handle staple for Cy3_398nm rod_69 |
| *CTCCTATCTCCAATCACTCCT*CATAAATACAGCAGTTGAATAATGGAA | Handle staple for Cy3_398nm rod_70 |

| | |
|---|---|
| *CTCCTATCTCCAATCACTCCT*TTAAACAGGGATCAAAAAATTATTTGC | Handle staple for Cy3_398nm rod_71 |

Handle staples for Cy3 fluorophore in a 501-nm long DNA calibration rod. (The following handles were added along with those for the 398-nm long DNA calibration rod)

| **Sequence (5' to 3')** | **Name** |
|---|---|
| *CTCCTATCTCCAATCACTCCT*GAAACAGCTTCAGAAAATAACGGAATA | Handle staple for Cy3_501nm rod_1 |
| *CTCCTATCTCCAATCACTCCT*GAAGGGATTCAGGTCTAGTTACCAGAA | Handle staple for Cy3_501nm rod_2 |
| *CTCCTATCTCCAATCACTCCT*GGAATTTGGCAAAGCGTTTTTAAGAAA | Handle staple for Cy3_501nm rod_3 |
| *CTCCTATCTCCAATCACTCCT*ACGGGAACTTCGAGCTTGAGTTAAGCC | Handle staple for Cy3_501nm rod_4 |
| *CTCCTATCTCCAATCACTCCT*GCCAGTGCAACTCCAAGCGCTAATATC | Handle staple for Cy3_501nm rod_5 |
| *CTCCTATCTCCAATCACTCCT*GGTTTTCCTAATTGCTGAACACCCTGA | Handle staple for Cy3_501nm rod_6 |
| *CTCCTATCTCCAATCACTCCT*TGCTGCAACGGATGGCACAGGGAAGCG | Handle staple for Cy3_501nm rod_7 |
| *CTCCTATCTCCAATCACTCCT*TCGCTATTTGCTGTAGATAGCAGCCTT | Handle staple for Cy3_501nm rod_8 |
| *CTCCTATCTCCAATCACTCCT*ACTGTTGGTAAAGTACAAACGATTTTT | Handle staple for Cy3_501nm rod_9 |
| *CTCCTATCTCCAATCACTCCT*ATCAAAAAAGCTCTCAATTGCGTAGAT | Handle staple for Cy3_501nm rod_10 |
| *CTCCTATCTCCAATCACTCCT*TTAAGAGGTACAGCGCGAAACAATAAC | Handle staple for Cy3_501nm rod_11 |
| *CTCCTATCTCCAATCACTCCT*CGTTTTAAGGATAACCATACCAAGTTA | Handle staple for Cy3_501nm rod_12 |
| *CTCCTATCTCCAATCACTCCT*CGGAAGCACAAGCTTTTATTCATTTCA | Handle staple for Cy3_501nm rod_13 |
| *CTCCTATCTCCAATCACTCCT*AGTACCTTCAGTCACGGATGAAACAAA | Handle staple for Cy3_501nm rod_14 |
| *CTCCTATCTCCAATCACTCCT*CATTTTTGGGCGATTATACATTTAACA | Handle staple for Cy3_501nm rod_15 |
| *CTCCTATCTCCAATCACTCCT*GAATATAAACGCCAGCTTAATGGAAAC | Handle staple for Cy3_501nm rod_16 |
| *CTCCTATCTCCAATCACTCCT*TATGCAACGAAGGGCGGAGTGAATAAC | Handle staple for Cy3_501nm rod_17 |

Handle staples for Cy3 fluorophore in 599 nm DNA calibration rod. (The following handles were added along with those for the 398-nm and 501-nm long DNA calibration rod)

| **Sequence (5' to 3')** | **Name** |
|---|---|
| *CTCCTATCTCCAATCACTCCT*GGCAAAGCTAACAGTTACAGCCATATT | Handle staple for Cy3_599nm rod_1 |
| *CTCCTATCTCCAATCACTCCT*CTTTCCGGGATTTAGTTTCCAGAGCCT | Handle staple for Cy3_599nm rod_2 |
| *CTCCTATCTCCAATCACTCCT*TATCGGCCATGGTCAAATCCTGAATCT | Handle staple for Cy3_599nm rod_3 |
| *CTCCTATCTCCAATCACTCCT*GTGCATCTTTGGGGCGTTAGTTGCTAT | Handle staple for Cy3_599nm rod_4 |
| *CTCCTATCTCCAATCACTCCT*GTCACGTTCTACTAATCTTGCGGGAGG | Handle staple for Cy3_599nm rod_5 |
| *CTCCTATCTCCAATCACTCCT*GAACAAACATCATACAAAGAACGCGAG | Handle staple for Cy3_599nm rod_6 |
| *CTCCTATCTCCAATCACTCCT*GAGCGAGTTAAGCAATAAATCAGATAT | Handle staple for Cy3_599nm rod_7 |
| *CTCCTATCTCCAATCACTCCT*TCCTGTAGTCGGTTGTCGTAGGAATCA | Handle staple for Cy3_599nm rod_8 |
| *CTCCTATCTCCAATCACTCCT*GAACGCCATACTTTTGATCGAGAACAA | Handle staple for Cy3_599nm rod_9 |
| *CTCCTATCTCCAATCACTCCT*ATTCCATAGCCATTCGTATTAATTAAT | Handle staple for Cy3_599nm rod_10 |
| *CTCCTATCTCCAATCACTCCT*AACGAGTACACCGCTTACATAGCGATA | Handle staple for Cy3_599nm rod_11 |
| *CTCCTATCTCCAATCACTCCT*TTTCGCAATCAGGAAGAGAGTCAATAG | Handle staple for Cy3_599nm rod_12 |
| *CTCCTATCTCCAATCACTCCT*ATTTTCATGCCAGTTTTCTGAGAGACT | Handle staple for Cy3_599nm rod_13 |
| *CTCCTATCTCCAATCACTCCT*CATCAATTGGTGTAGAGGTTGGGTTAT | Handle staple for Cy3_599nm rod_14 |
| *CTCCTATCTCCAATCACTCCT*TCCAATAAGGCGGATTTGCAAATCCAA | Handle staple for Cy3_599nm rod_15 |
| *CTCCTATCTCCAATCACTCCT*AGCAAAATAACAACCCAAAACTTTTTC | Handle staple for Cy3_599nm rod_16 |
| *CTCCTATCTCCAATCACTCCT*AAGCTAAACCAGCTTTATCTTCTGACC | Handle staple for Cy3_599nm rod_17 |
| *CTCCTATCTCCAATCACTCCT*CCCTGTAATCAAAAATCCGACCGTGTG | Handle staple for Cy3_599nm rod_18 |

Handle staples for Cy3 fluorophore in 658 nm DNA calibration rod. (The following handles were added along with those for the 398-nm, 501-nm, and 599-nm long DNA calibration rod)

| **Sequence (5' to 3')** | **Name** |
|---|---|
| *CTCCTATCTCCAATCACTCCT*TTTGTTAAAAGGATAACAAGAACGGGT | Handle staple for Cy3_658nm rod_1 |
| *CTCCTATCTCCAATCACTCCT*CGTTAATATTAAATGCAATCAATAATC | Handle staple for Cy3_658nm rod_2 |
| *CTCCTATCTCCAATCACTCCT*AAGATTGTAGATTCAAATCCCATCCTA | Handle staple for Cy3_658nm rod_3 |
| *CTCCTATCTCCAATCACTCCT*CGGTTGATTCAAATCAACAATAGATAA | Handle staple for Cy3_658nm rod_4 |
| *CTCCTATCTCCAATCACTCCT*TCGTAAAACTAGCTGAGTTCAGCTAAT | Handle staple for Cy3_658nm rod_5 |
| *CTCCTATCTCCAATCACTCCT*TTCAACGCATCAGCTCAATAAACACCG | Handle staple for Cy3_658nm rod_6 |
| *CTCCTATCTCCAATCACTCCT*CATATATTTTTGTTAGCCTGTTTAGT | Handle staple for Cy3_658nm rod_7 |
| *CTCCTATCTCCAATCACTCCT*GTAGGTAAATAAGCAACTTACCAGTAT | Handle staple for Cy3_658nm rod_8 |
| *CTCCTATCTCCAATCACTCCT*GGAGACAGAATCAGAAGGCTTAATTGA | Handle staple for Cy3_658nm rod_9 |
| *CTCCTATCTCCAATCACTCCT*AGGGTAGCAACAAGAGGCCAGTAATAA | Handle staple for Cy3_658nm rod_10 |

**Table S7**

Antihandles for DNA calibration rod.

| **Sequence (5' to 3')** | **Name** |
|---|---|
| biotin/TGGATATGGTGGAGAGGAGAG | Biotin-labeled antihandle |
| Cy3/AGGAGTGATTGGAGATAGGAG | Cy3-labeled antihandle |
